# A brain-based universal measure of attention: predicting task-general and task-specific attention performance and their underlying neural mechanisms from task and resting state fMRI

**DOI:** 10.1101/2021.02.13.431091

**Authors:** Kwangsun Yoo, Monica D Rosenberg, Young Hye Kwon, Emily W Avery, Qi Lin, Dustin Scheinost, R Todd Constable, Marvin M Chun

## Abstract

Attention is central for many aspects of cognitive performance, but there is no singular measure of a person’s overall attentional functioning across tasks. To develop a universal measure that integrates multiple components of attention, we collected data from more than 90 participants performing three different attention-demanding tasks during fMRI. We constructed a suite of whole-brain models that can predict a profile of multiple attentional components – sustained attention, divided attention and tracking, and working memory capacity – from a single fMRI scan type within novel individuals. Multiple brain regions across the frontoparietal, salience, and subcortical networks drive accurate predictions, supporting a universal (general) attention factor across tasks, which can be distinguished from task-specific attention factors and their neural mechanisms. Furthermore, connectome-to-connectome transformation modeling enhanced predictions of an individual’s attention-task connectomes and behavioral performance from their rest connectomes. These models were integrated to produce a new universal attention measure that generalizes best across multiple, independent datasets, and which should have broad utility for both research and clinical applications.

## Introduction

Attention has a ubiquitous role in perception and cognition (Chun et al., 2011). We are endlessly exposed to all kinds of overflowing sensory information, and the ability to deploy attention over space and to sustain it over time is crucial in everyday life. We have, however, limited cognitive capacity, and therefore must selectively process information most relevant to our actions. Attentiveness explains behavioral performance fluctuations both within and across individuals (Weissman et al., 2006), and attention deficits are common in mental illness and symptomatic of brain damage (Biederman et al., 1991; Heinrichs and Zakzanis, 1998; Levin et al., 1987).

Despite this central importance of attention, clinicians and researchers lack a standardized way to measure a person’s overall attentional functioning. Although no mental process can be reduced to a single number, both research and clinical practice can benefit from having standardized and quantifiable measures to facilitate comparison across and within individuals (Rosenberg et al., 2016a, 2018). For example, intelligence research and education practice benefits from the ability to measure g, as an index of fluid intelligence (Deary et al., 2010). A comparable index is lacking for attention, despite its pervasive role in modulating most perceptual and cognitive processes.

The fact that there are so many different tasks to examine attention functioning reflects that attention is not a unitary construct but rather multifaceted (Chun et al., 2011). People’s attentional abilities may vary along the multiple dimensions of attention. These differences in attention functions among individuals can be measured by extensive behavioral tasks; however, an overload of tasks not only makes subjects feel fatigued, which may affect task performance, but also requires a substantial period of time. Therefore, it is important to understand what is common and what is unique among the different attention tasks and to try to predict them with minimal testing.

The literature lacks a systematic investigation of the general and specific factors of attentional processes and the underlying neural architectures supporting general and specific aspects of attention across an array of tasks. One behavioral study examined a set of cognitive tasks known to employ executive functions, including attention and working memory. This study showed that the nine tested tasks are not completely independent but share common and separable components (Miyake et al., 2000). Another study also revealed a general behavioral factor that is shared by multiple attention task paradigms in common as well as specialized factors that are unique to specific tasks (Huang et al., 2012). For the nine primary and eight secondary tasks involved in the study, one common component explained substantial variance in performance across most of the tasks. However, this general attention factor could not be derived from an individual task, and the behavioral study did not explore the underlying neural mechanisms across tasks. The frontal and parietal cortices are well-known to control attention (Corbetta and Shulman, 2002; Kanwisher and Wojciulik, 2000), but most studies have not compared their engagement across multiple attention tasks. In addition, only recently have studies begun to predict individual attentional behaviors from brain scans (Kucyi et al., 2017; Rosenberg et al., 2016a, 2017; Wu et al., 2020).

Here we seek to develop a universal attention profile measure that can quantify a person’s performance across the different cognitive demands of sustained attention, divided attention and tracking, and working memory. The attention profile includes task-specific measures and a universal measure that generalizes across tasks. We use a neuroimaging-based data-driven approach called connectome-based predictive modeling (CPM; Shen et al., 2017) that develops computational models to accurately predict an unseen, novel individual’s trait and behavior from their brain activity. This is based on the whole-brain pattern of functional connectivity (synchronized fluctuation of time-series signals from distributed brain regions), which is unique to each individual as a fingerprint and predictive of their behaviors (Cohen and D’Esposito, 2016; Finn et al., 2015; Gratton et al., 2020; Woo et al., 2017). CPM accurately predicts a variety of individual behaviors and traits, including intelligence (Finn et al., 2015; Yoo et al., 2019), attention (Rosenberg et al., 2016a, 2016b, 2018; Yoo et al., 2018), memory (Avery et al., 2019; Lin et al., 2018; Zhang et al., 2020), language (Tomasi and Volkow, 2020), creativity (Beaty et al., 2018) and personalities (Cai et al., 2020; Hsu et al., 2018; Jiang et al., 2018).

One of the earliest CPM studies introduced a model to predict an individual’s ability to sustain attention (Rosenberg et al., 2016a). The study demonstrated that the brain’s functional organization is predictive of behavioral performance in the gradual-onset continuous performance task (gradCPT, Esterman et al., 2013). In addition, the CPM of sustained attention generalized to predict individual performance in a stop-signal task, performances in the Attention Network Task (ANT; Fan et al., 2005) and symptom severity in patients with attention-deficit/hyperactivity disorder (Rosenberg et al., 2016a, 2016b, 2018). Sustained attention is, however, just one aspect of human attention (Chun et al., 2011).

For a more comprehensive assessment of attention, we collected original behavioral and fMRI data from more than 90 participants performing three different attention tasks during fMRI scanning. The three tasks include the gradCPT to measure sustained attention, multiple object tracking (MOT) to measure divided attention and tracking, and a visual short-term memory (VSTM) task to assess working memory capacity as a form of internal attention (Chun et al., 2011; Engle, 2002).

The primary goal of this study is to develop a battery of whole-brain functional network models that can measure how individuals vary in their attentional abilities. We can predict task-specific attention measures, or we can unify the models to generate a universal attention measure. The suite of attention models developed in this study accurately predicts individual behaviors from functional connectivity measured during task performance as well as from the resting state. A quantitative profile from a single fMRI session facilitates comparison across individuals and between sites, and it enables longitudinal tracking of individuals (Rosenberg et al., 2020) to assess development, aging, and intervention. Moreover, we leveraged the network models to probe brain systems that support common and separable factors for attention functions measured during the tasks. The investigation of functional anatomy across attentional functions sheds light on the relationships between components of attention and the brain networks that support them.

Lastly, we developed new ways to significantly improve how patterns of brain networks supporting multiple attentional processes can be drawn from resting-state data alone. This is essential because attention tasks vary widely and are difficult to standardize across studies and settings. It is also impractical to ask participants, especially patients or children, to engage in many different attention tasks, especially inside a brain scanner. If a profile of attention measures can be derived from resting-state data, it would have significant utility for both research applications and clinical practice. To enhance predictability from resting-state data, we utilized a novel method called connectome-to-connectome (C2C) state transformation modeling. The C2C framework generates individual task-related connectomes from their rest connectomes with high specificity and improved behavioral prediction performance (Yoo et al., 2020).

## Results

### Individual behaviors in three attentional tasks

We first examined the reliability and similarity of individual performance in the three attention tasks. To address the reliability of the behavioral measure of each task, we assessed the behavioral similarity between two sessions using Pearson’s correlation. Among the total of 94 subjects analyzed in this study, 65, 69, and 72 subjects had available behavioral performances in both sessions of gradCPT, MOT, and VSTM, respectively. The individual behaviors were significantly correlated between two sessions in all three tasks (gradCPT: *r*=0.738, *p*=2.42×10^−12^; MOT: *r*=0.625, *p*=9.40×10^−9^; VSTM: *r*=0.577, *p*=1.16×10^−7^; significant under Bonferroni corrected *p*<0.05, **Supplementary Figure S1**). These significant correlations between sessions indicate that individual behavioral performances were reliably measured in all tasks.

We then tested how similarly participants performed in the three attention tasks. To assess the similarity of individual behaviors between the three different tasks, we estimated Pearson’s correlation of individual performances between them. Individual performances were positively correlated between every pair of three attention tasks (gradCPT-MOT: *r*=0.366, *p*=2.86×10^−4^; gradCPT-VSTM: *r*=0.479, *p*=1.04×10^−6^; MOT-VSTM: *r*=0.459, *p*=3.35×10^−6^; significant under Bonferroni corrected *p*<0.05, **Supplementary Figure S2**).

### CPMs of three attentional tasks

We built a battery of predictive models of attentional functions (**Table 1**). We constructed nine CPMs of attention that differed in cognitive states of fMRI scans (performing attention tasks, resting-state, or movie-watching) and target attention tasks (gradCPT, MOT, and VSTM). In detail, three models were trained to predict individual gradCPT performance, another three were to predict MOT performance, and the other three were to predict VSTM performance. Among the three models of each task, one model was trained using the corresponding task fMRI, another one was trained using rest fMRI, and the other one was trained using movie fMRI. We evaluated these nine CPMs in predicting an unseen individual’s task performance using LOOCV. We assessed the model’s prediction accuracy by correlating model-predicted and observed behavioral scores. In all three tasks, predicted behavioral scores significantly correlated with actual task scores (*p*s<0.05 FWE-corrected using permutation) when CPMs were trained using task fMRI (the *top row in* **Figure 1**). A significant positive correlation indicates that CPMs accurately predict individual differences in task performance. The CPMs trained using movie fMRI also accurately predicted individual performance in all three tasks (*p*s<0.05 FWE-corrected using permutation, *bottom row* in **Figure 1**). In contrast, rest fMRI-based models only predicted behavioral scores in gradCPT, and failed to predict scores in MOT and VSTM (the *middle row* in **Figure 1**).

**Table 1.**
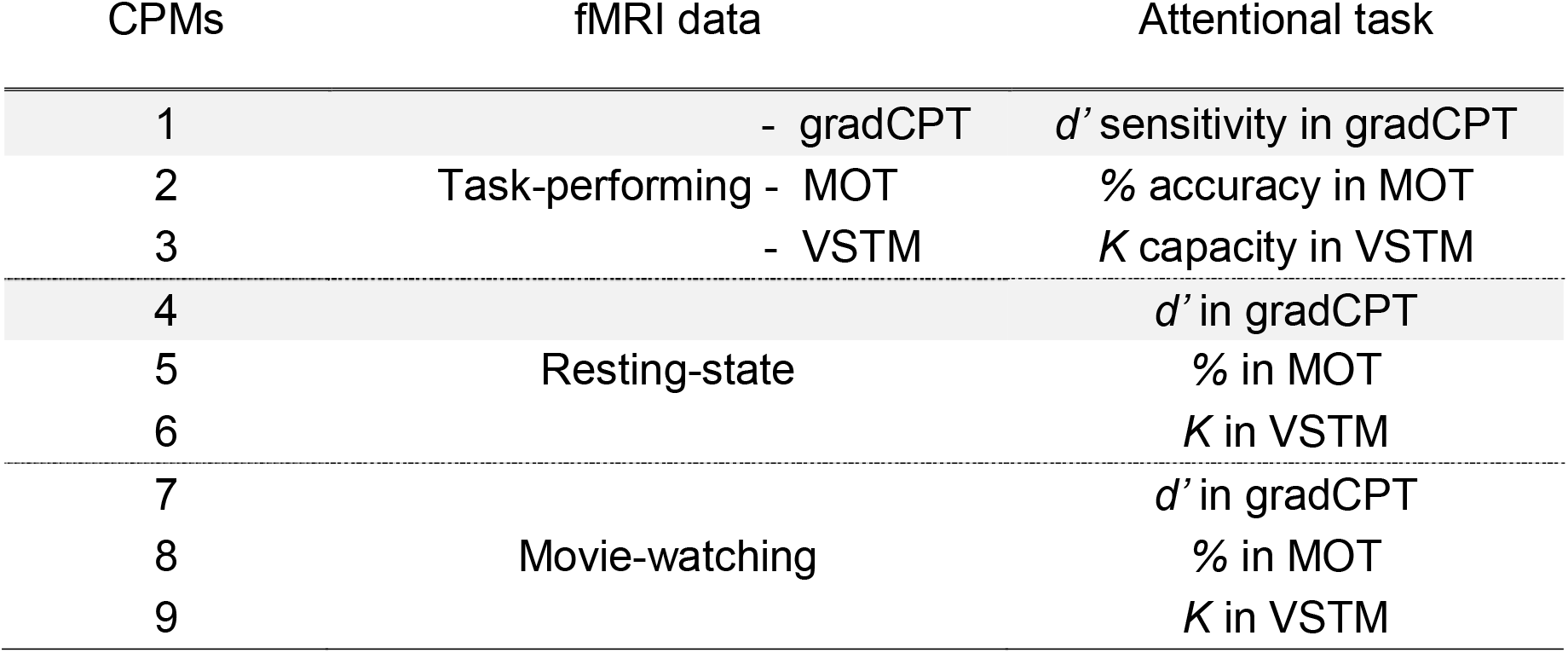
Nine CPMs of attention. Two models in gray shade are replications of our previous study (Rosenberg et al., 2016a), but with different samples. The other seven models were newly developed in the current study. GradCPT: gradual-onset continuous performance task, MOT: multiple object tracking, and VSTM: visual short-term memory.

**Figure 1.**
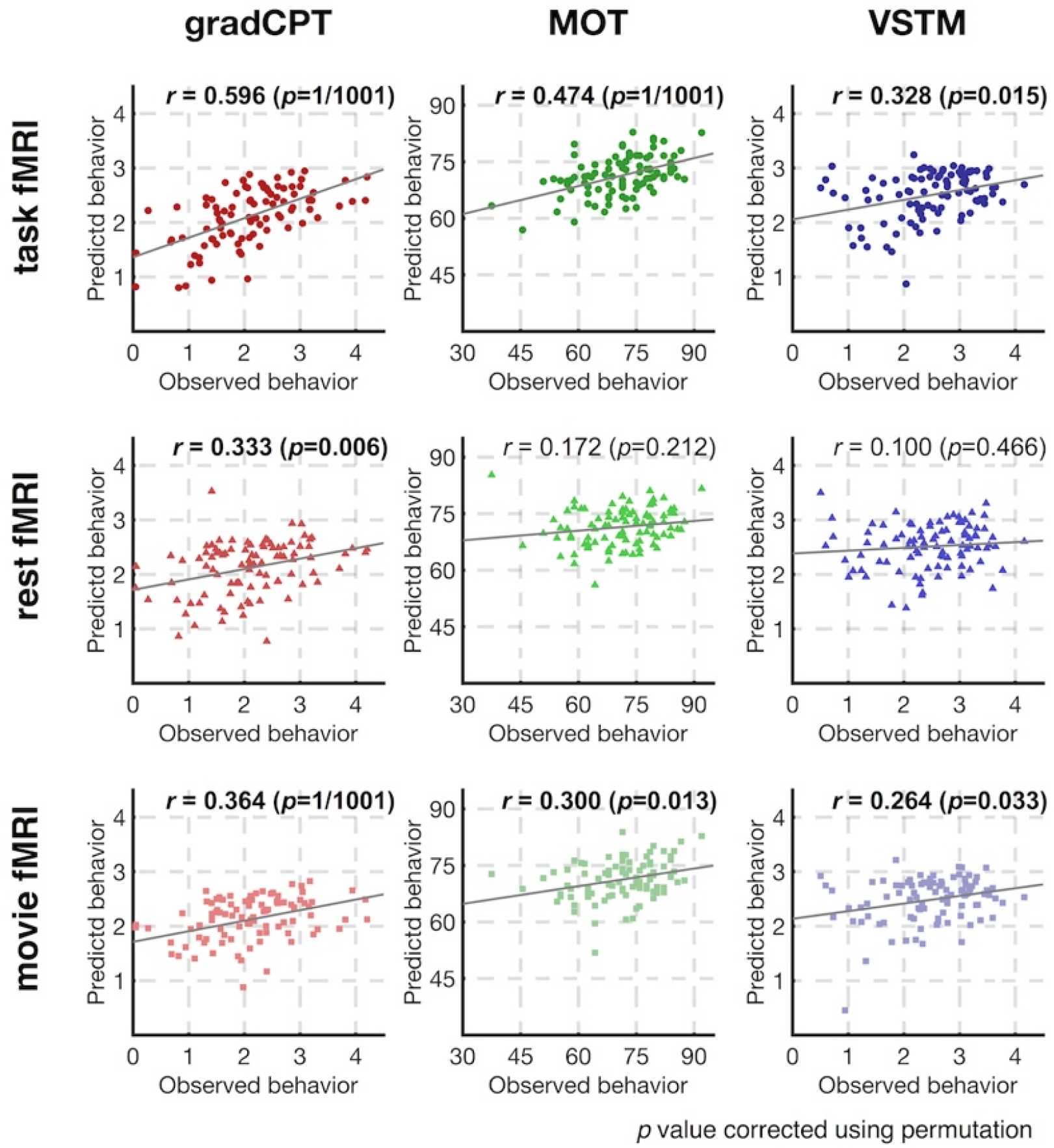
Prediction accuracy of nine CPMs. Rows represent the fMRI data used in model training and prediction, and columns represent the target attention task. Models’ prediction accuracies were assessed by correlating model-predicted behavioral scores and observed scores. *P* values were obtained using 1,000 permutations and corrected for multiple tests (three correlation tests with task fMRI-based models, three tests with rest fMRI-based models, and three tests with movie fMRI-based models). GradCPT: gradual-onset continuous performance task, MOT: multiple object tracking, and VSTM: visual short-term memory.

In addition to examining correlations, we also calculated prediction q^2^ (Scheinost et al., 2019). The prediction q^2^ is a normalized version of mean square error, assessing the model’s numerical accuracy in predicting individuals’ actual behavioral scores relative to simply guessing their population mean. Thereby, this prediction q^2^ assessment complements the correlation-based evaluation which measures the model accuracy in predicting individual differences in behavior. The prediction q^2^ result corroborated the successful prediction of attention functions (**Supplementary Figure S3**). Three models using task fMRI data successfully predicted individual behavioral scores (*p*s<0.05 FWE-corrected using permutation). The models using movie fMRI data also accurately predicted individual scores (*p*s<0.05 FWE-corrected using permutation). The rest fMRI-based model predicted only behavioral scores in gradCPT.

### CPMs generalize to predict individual behaviors in different tasks

We investigated whether the CPM of each task generalizes to different attention tasks and different cognitive states of fMRI. The CPMs trained using task fMRI successfully generalized to different attention tasks (*the top left 3 by 3 subpart* in **Figure 2**, *p*s<0.05 FWE corrected using permutation). Interestingly, all three task-based models predicted individual performances in gradCPT better than their own corresponding tasks (**Figure 2** and **Supplementary Figure S4**). For example, the CPM trained using MOT fMRI predicted unseen subjects’ gradCPT performance better than MOT performance. In addition, the current results showed that movie fMRI (*the bottom right 3 by 3 subpart* in **Figure 2**) better predicts individual attention behaviors compared to rest fMRI (*the center 3 by 3 subpart* in **Figure 2**). The models with movie fMRI provided successful predictions in general (four successful predictions among six cross-task prediction cases, *p*s<0.05 FWE corrected using permutation), whereas the models with rest fMRI could predict behaviors only in gradCPT.

**Figure 2.**
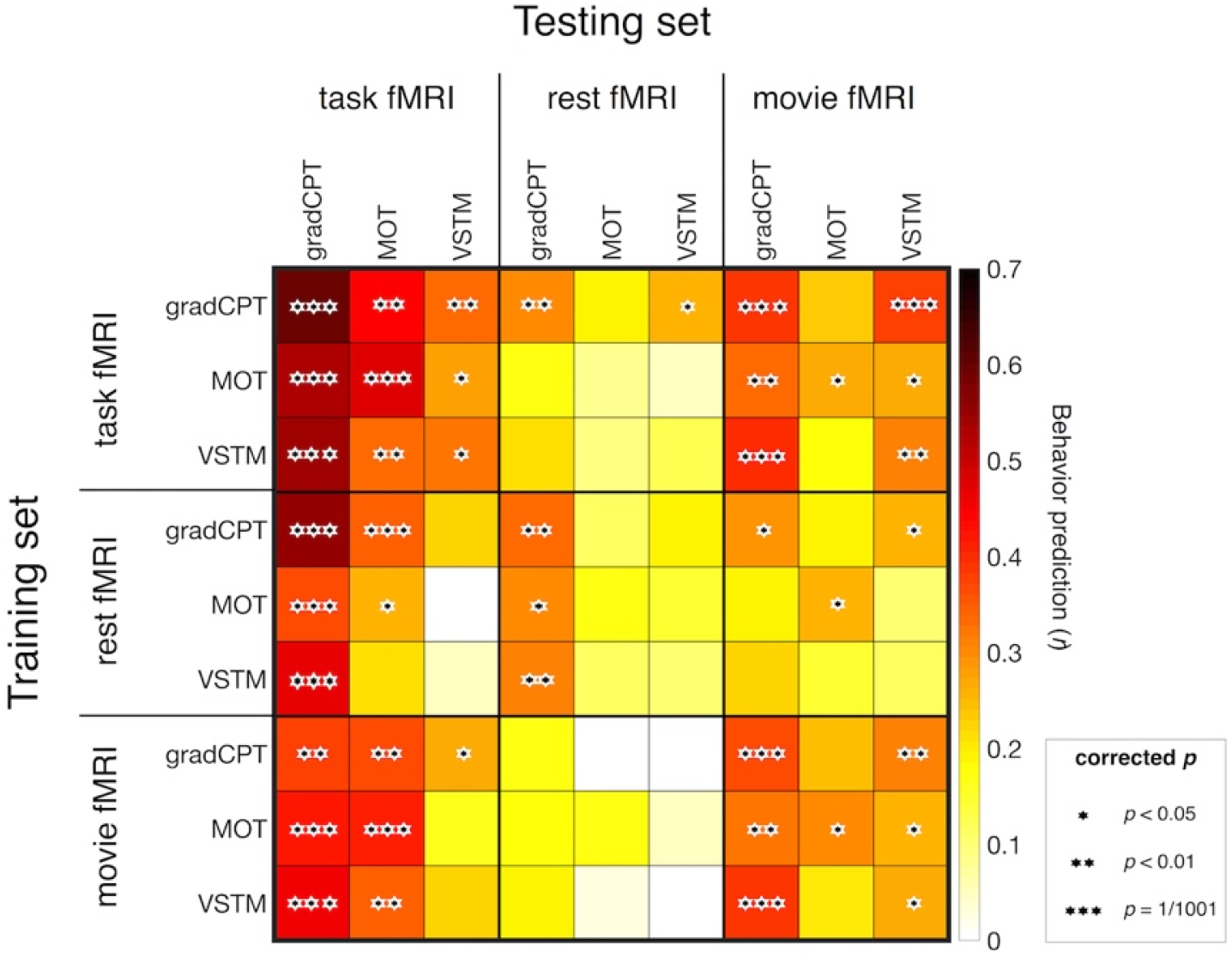
Cross-prediction results across all cognitive states and attention tasks (*: *p*<0.05, **: *p*<0.01, and ***: *p*=1/1,001, corrected using permutation). Rows represent combinations of fMRI data and behavior scores used in model construction, and columns represent combinations of fMRI data and behavior scores used in model validation. On-diagonal elements represent the nine within-task prediction results and off-diagonal elements represent the cross-task predictions. Models’ prediction accuracies were assessed by correlating model-predicted and observed behavioral scores. *P* value was obtained using 1,000 permutations and corrected for multiple tests. The models with task fMRI successfully generalized to different attention tasks (*the top left 3 by 3 subpart*), and the models with movie fMRI also generalized to different tasks to lesser degrees (*the bottom right 3 by 3 subpart*). GradCPT: gradual-onset continuous performance task, MOT: multiple object tracking, and VSTM: visual short-term memory.

### Generalizability of CPMs is not driven by correlated behaviors between tasks

We employed two different approaches to examine whether the generalizability of CPMs between tasks is fully driven by the correlations in behavior between them. First, we constructed variants of the original nine CPMs with connectivity features controlled for the performance in the two non-target tasks. Edges that are correlated with performance in a target task were first selected as features, and amongst these, edges that correlated with performance in either of the other two tasks were excluded. Hence, these new models include predictive edges that correlated only with individual behavioral performance in a target task, but not with behaviors in any of the other two transfer tasks; the resulting models contained only task-specific edges. All other steps except this feature selection remained the same as the original model construction. We then assessed the generalizability of these nine variant models. This analysis yielded a pattern of successful cross-task prediction similar to the original models (**Supplementary Figure S5**). The models with task fMRI successfully predicted behaviors in different tasks (*p*s<0.05 FWE corrected using permutation). Predictions by models with movie fMRI in general were also still successful (three among six cross-task predictions, *the bottom left 3 by 3 subpart* in **Supplementary Figure S5**, *p*s<0.05 FWE corrected using permutation). This result indicates that the generalizability of the attention CPMs between different tasks is not dependent on the correlated behaviors and the shared connectivity features across different tasks.

As a second way to rule out the effect of correlated behavior in generalizing the models, we re-assessed the prediction accuracy of the original nine CPMs while controlling for performance in the non-target tasks. In this case, we did not construct new models. Instead, we used partial correlations to evaluate the generalizability of the original models. We correlated model-predicted and observed behavior performance with controlling for performance in non-target tasks. For example, to examine if a model trained to predict gradCPT scores generalizes to predict MOT, we calculated the partial correlation between model-predicted scores and observed MOT scores with controlling for observed gradCPT scores. The partial correlation results were consistent with the results from the models with task-specific connectivity (**Supplementary Figure S6**, *p*s<0.05). The results further support that predictive models generalize well across different attention tasks even when controlling for correlations between behavior.

### Predictive anatomy of the original CPMs of attentional tasks

We explored the predictive anatomy of the original CPMs of three attention tasks. The distribution of predictive connectivity is similar across three tasks on a macro scale (**Figure 3A**). In all three models, predictive connectivity is distributed across the whole brain, coinciding with the previous reports (Rosenberg et al., 2016a). The salience, visual II, frontoparietal, subcortical, and cerebellum networks were shown to play major roles in the CPMs (*darker color* in **Figure 3A**). To test whether there is a general connectivity component underlying the three attention tasks, we tracked the overlap of predictive connectivity between three task-based models. (*the bottom right* in **Figure 3A**). The connectivity of the salience, subcortical, and cerebellum networks were commonly involved in all three task-based CPMs for both positive and negative networks. In addition, the negative network also included connectivity of the frontoparietal and visual II networks.

**Figure 3.**
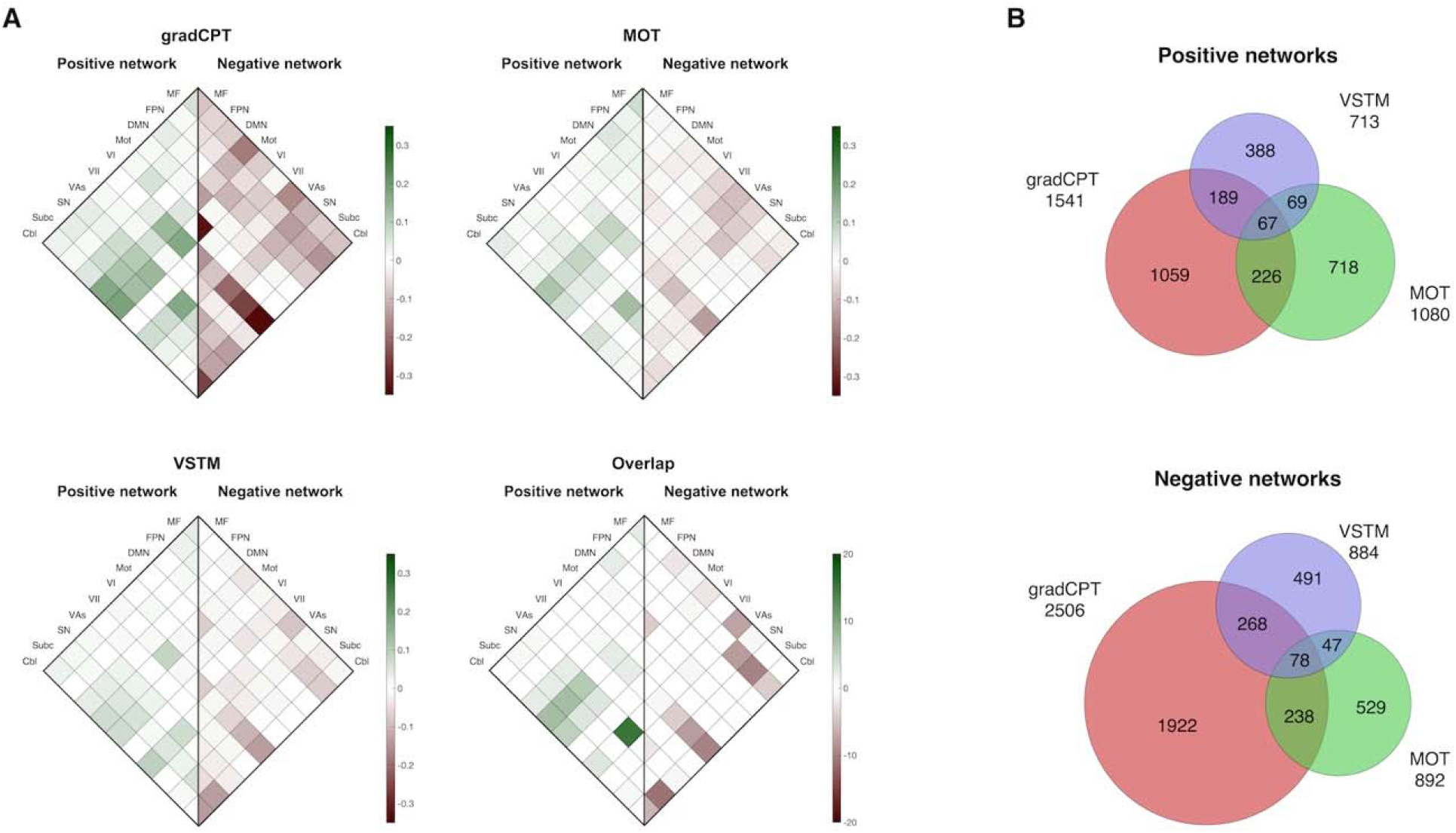
**A**. Predictive anatomy of three task-based CPMs. The scale bar in gradCPT, MOT and VSTM represents the relative ratio of predictive functional connections to all possible number of functional connections between networks with a sign representing whether the functional connection is in a positive or negative network. The scale bar in overlap represents the actual number of predictive functional connections with a sign representing whether the connectivity is in a positive or negative network. GradCPT: gradual-onset continuous performance task, MOT: multiple object tracking, and VSTM: visual short-term memory. MF: medial-frontal network, FPN: frontoparietal network, DMN: default mode network, Mot: motor, VI: visual I, VII: visual II, VAs: visual associations, SN: salience network, Subc: subcortex, Cbl: cerebellum. **B**. The number of predictive functional connections of three task-based CPMs in positive and negative networks.

### CPMs are robust across the number of predictive connectivity features

Although we found CPMs of three tasks generalize well between tasks, predictive performance was numerically different across them (**Figure 1 & 2**). The number of predictive connectivity features was different across the models; for gradCPT there were 1,541 and 2,506 edges in the positive and negative networks, respectively, whereas for VSTM there were 713 and 884 edges, less than half the number in gradCPT (**Figure 3B**). The numbers of predictive connectivity features for MOT were in-between gradCPT and VSTM, 1,080 and 892 edges in the positive and negative networks, respectively. We, therefore, examined if the better performance of gradCPT model is due to the larger number of predictive connectivity features compared to that of MOT and VSTM. Here, we constructed variants of the nine models with different feature selection criteria. In our original model, we selected edges that were significantly correlated with individual behaviors of interest at *p*<0.05. In the current analysis, we selected the top *n* edges whose correlations with behaviors (either positive or negative) are most significant. In this way, we control the number of predictive edges to be the same between different models. We observed that model predictions remain similar with this new feature selection criteria (**Supplementary Figure S7**). Across the range of selection thresholds, the gradCPT CPM predicted most accurately as the original CPM model. This analysis indicates that the size of predictive network is not a driving factor for model performances and that CPM modeling is robust against the choice of feature selection threshold.

### Head motion during attention tasks cannot explain the accurate prediction of CPMs

To test whether there is any relationship between head motion and task performance, we correlated head motion, measured by mean FD, and task performance across individuals. MOT performance was negatively correlated with head motion during all three tasks, rest, and movie scans (*p*<0.05 Bonferroni corrected, **Supplementary Table S1**). Performance in gradCPT and VSTM were not correlated with head motion.

Since head motion was correlated with task performance in MOT, we reevaluated the performance of the original CPMs to confirm that the prediction of behavioral performance in MOT was not biased by head motion, measured by mean FD. To account for head motion in behavior prediction, we assessed the prediction accuracy of models by running partial correlations between the model-predicted and observed behavioral scores while controlling for head motion during the observed behavior. For example, when we trained task-based model of gradCPT and tested this model in predicting the performance in MOT task, we ran a partial correlation between the predicted scores and the observed MOT scores while controlling for head motion during the MOT task. Prediction performance of the models remained mostly significant even after controlling for head motion (**Supplementary Figure S8**). Specifically, the prediction for MOT performance showed the largest drop in accuracy compared to that when not controlled for head motion (no significance, *ps*>0.07 in **Supplementary Figure S8B**). Considering that only the performance in MOT task was correlated with head motion, this modulation in predicting MOT is not surprising. Overall, the present analyses demonstrated that the successful prediction of the constructed models was not based on head motion and that the CPMs established a reliable association between functional brain connectivity and attentional behaviors.

### CPMs of a universal (general) attention factor accurately predict individual behaviors in three tasks

To investigate if we can build a successful prediction model with a universal attention factor, we performed different variations of predictive modeling. We trained nine predictive models to utilize edges that were correlated with all three task behaviors as features; the models used only connectivity features shared across tasks. It is worth noting that these new models have significantly fewer predictive edges compared to the original models (**Supplementary Table S2**). However, these models’ prediction accuracy and generalizability were almost identical with the nine original models. The models with task fMRI fully generalized to different attention tasks (*p*s<0.05 FWE corrected using permutation, **Figure 4A**). The predictive connectivity features of task fMRI-based models were distributed among multiple brain networks, mainly in the subcortical, cerebellum, frontoparietal, motor, visual II and salience networks (**Figure 4B**). The prediction accuracy with all task-related edges exhibited similar patterns to those of the original models, and the generalizability was not significantly improved (**Supplementary Figure S9**).

**Figure 4.**
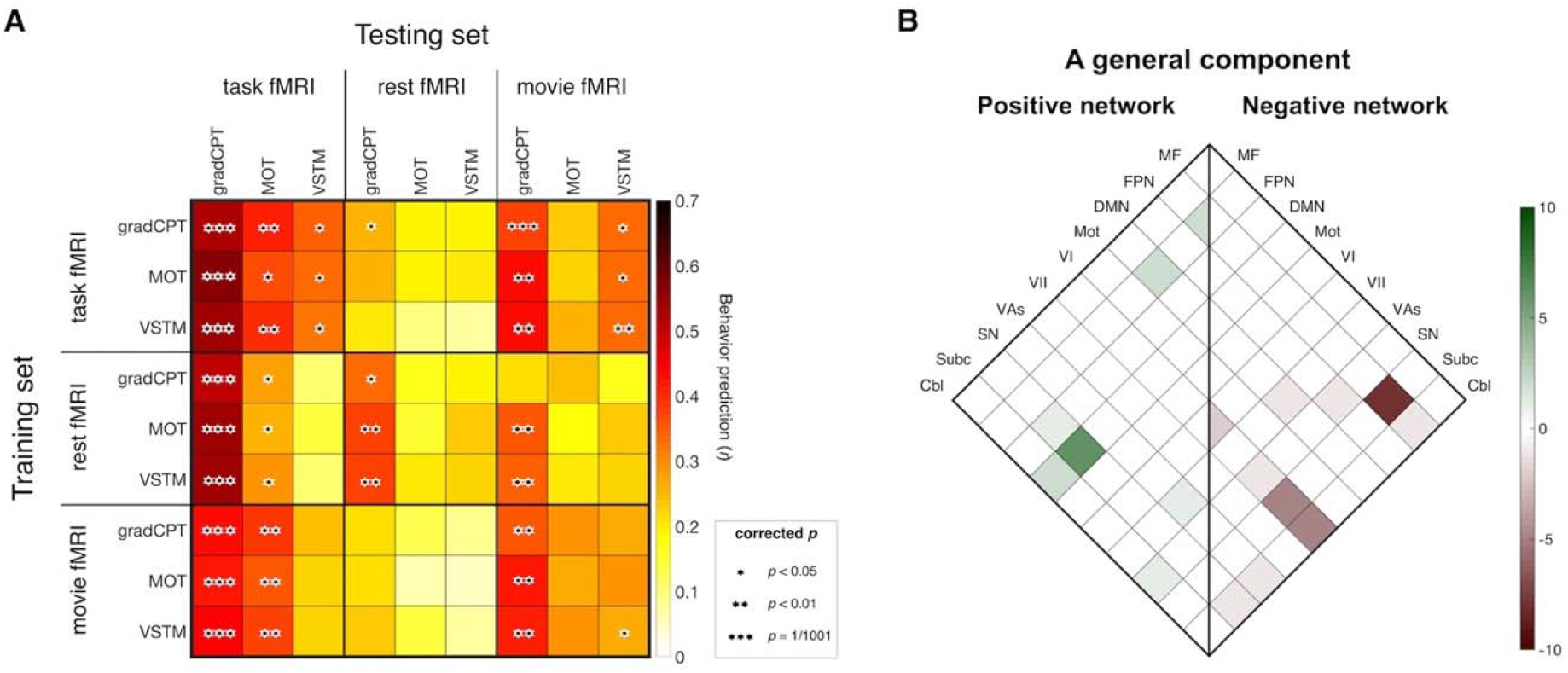
**A**. Cross-prediction results with a general component of three tasks (i.e., task-common edges; *: *p*<0.05, **: *p*<0.01, and ***: *p*=1/1,001, corrected using permutation). The models with task fMRI successfully generalized to different attention tasks (*the top left 3 by 3 subpart*). **B**. Predictive functional connections of a universal attention component that are shared in all three task fMRI-based CPMs. The scale bar represents the actual number of selected edges with a sign representing whether the functional connection is in a positive or negative network. GradCPT: gradual-onset continuous performance task, MOT: multiple object tracking, and VSTM: visual short-term memory. MF: medial-frontal network, FPN: frontoparietal network, DMN: default mode network, Mot: motor, VI: visual I, VII: visual II, VAs: visual associations, SN: salience network, Subc: subcortex, Cbl: cerebellum.

### Multiple brain networks contribute to prediction of attentional behaviors

We scrutinized the brain networks driving accurate behavior predictions. We defined ten networks (Finn et al., 2015; Noble et al., 2017), and to measure their contributions to performance, we computationally lesioned all the nodes in each network. Each of the ten networks was lesioned iteratively. After lesioning each network, we trained and tested three task fMRI-based CPMs in the same way the original three task-based models were constructed. We found that lesioning one network did not significantly decrease predictive models’ accuracy (**Figure 5A**). The three within-task predictions remained significant for all networks (*p*s<0.05 FWE corrected using permutation). All cross-task prediction cases also remained significant for all networks, except when lesioning the motor network and subcortical network for MOT to VSTM prediction (*p*s<0.05 FWE corrected using permutation).

**Figure 5.**
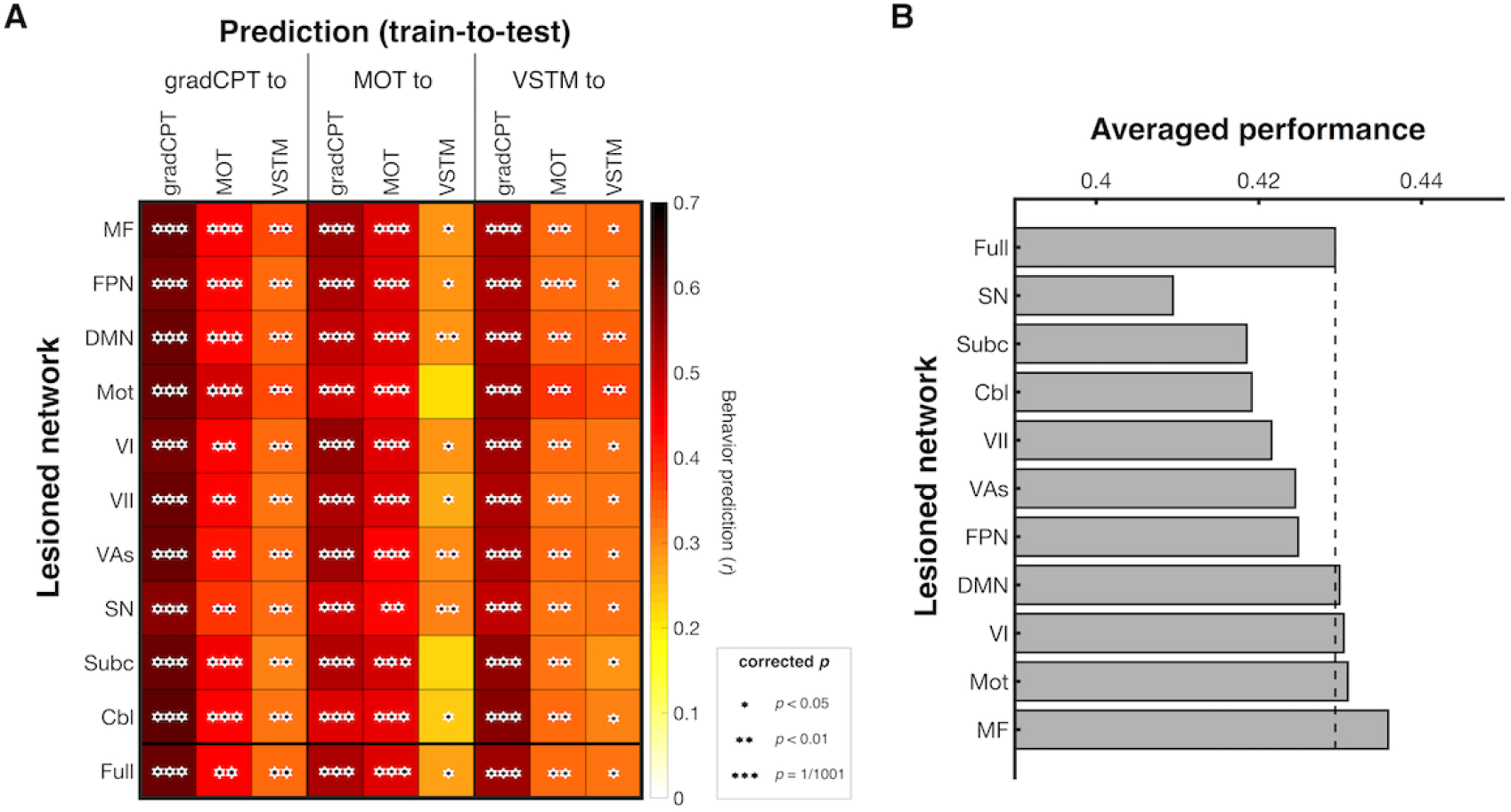
Behavioral prediction after lesioning each network. (*: *p*<0.05, **: *p*<0.01, and ***: *p*=1/1,001, corrected using permutation). **A**. Rows represent lesioned network and columns represent combinations of training and testing data. Models’ prediction accuracies were assessed by correlating model-predicted and observed behavioral scores. *P* value was obtained using 1,000 permutations and corrected for multiple tests. **B**. The average prediction performance of each network. This was obtained by averaging nine prediction cases in each row in **A**. GradCPT: gradual-onset continuous performance task, MOT: multiple object tracking, and VSTM: visual short-term memory. MF: medial-frontal network, FPN: frontoparietal network, DMN: default mode network, Mot: motor, VI: visual I, VII: visual II, VAs: visual associations, SN: salience network, Subc: subcortex, Cbl: cerebellum.

We further assessed the effect of lesioning each network on prediction (numerical decrease in accuracy) (**Figure 5B**). We found that the salience network, followed by the subcortical and cerebellum networks, is the most important in the attention prediction. To confirm that the lower performance was not driven by the smaller number of survived connections used in model training after lesioning, we tested the correlation between prediction performance and the number of survived connections (**Supplementary Table S3**). There was no positive correlation, indicating that the salience network (and the subcortical and cerebellum networks) indeed plays a major role in prediction of performance across attention tasks.

Next, we performed a complementary analysis to examine the predictive power of each network directly (**Figure 6A**). In this analysis, we built predictive models only using within-network connectivity of each network, iteratively for each network. This analysis confirmed that the subcortical, frontoparietal, and salience networks are the most predictive networks (**Figure 6B**). Although the prediction performance was lower than the original models’ accuracy, the prediction accuracy by each of these networks is notable given the significantly fewer number of features in these models.

**Figure 6.**
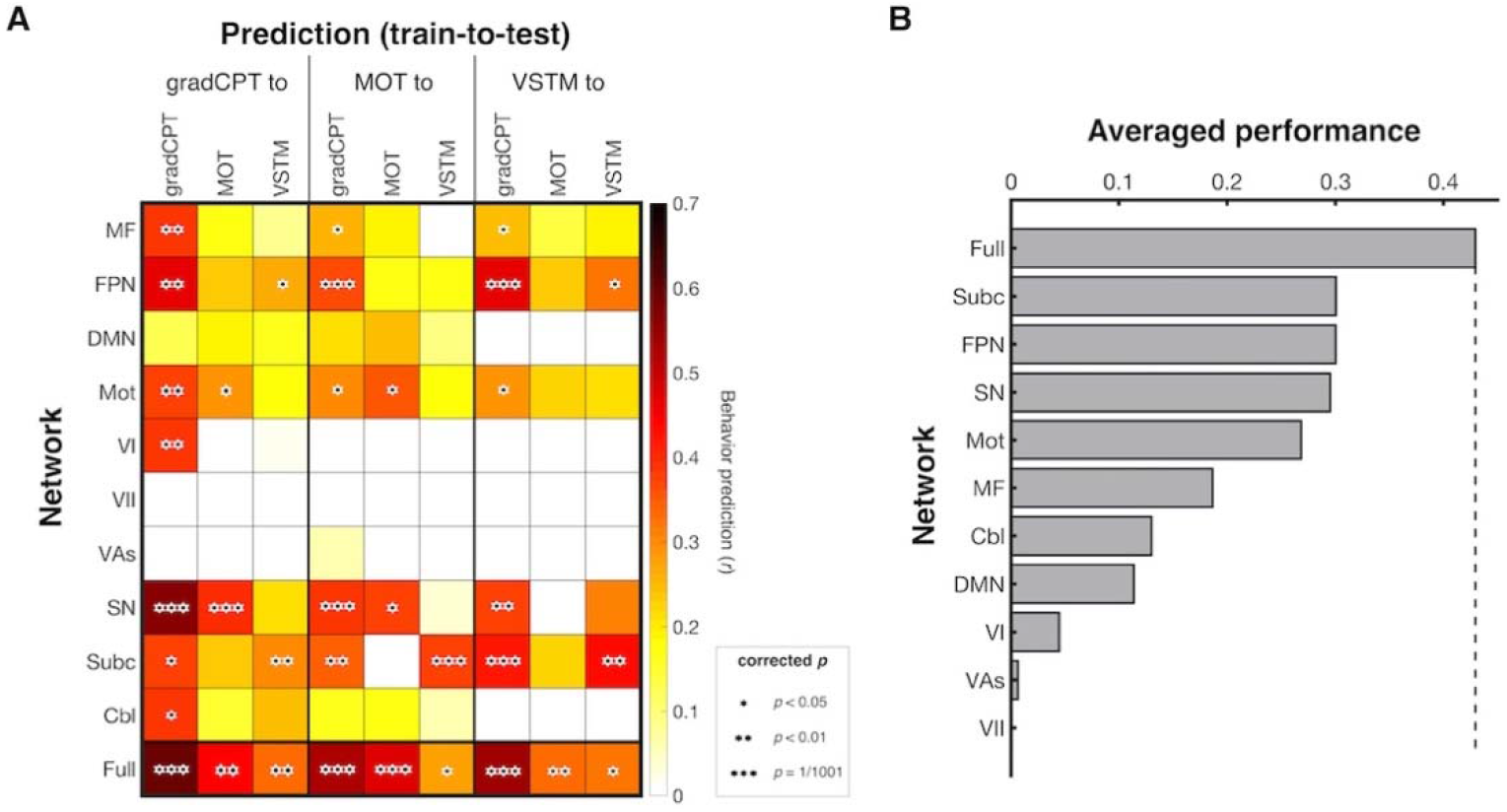
Cross-prediction using only within-network connectivity (*: *p*<0.05, **: *p*<0.01, and ***: *p*=1/1,001, corrected using permutation). **A**. Rows represent each canonical network and columns represent combinations of training and testing data. Models’ prediction accuracies were assessed by correlating model-predicted and observed behavioral scores. *P* value was obtained using 1,000 permutations and corrected for multiple tests. **B**. The average prediction performance of each network. This was obtained by averaging nine prediction cases in each row in **A**. GradCPT: gradual-onset continuous performance task, MOT: multiple object tracking, and VSTM: visual short-term memory. MF: medial-frontal network, FPN: frontoparietal network, DMN: default mode network, Mot: motor, VI: visual I, VII: visual II, VAs: visual associations, SN: salience network, Subc: subcortex, Cbl: cerebellum.

Note that this analysis examined the predictive power of only edges located within a network of interest; in contrast, the previous lesioning analysis examined the importance of edges within a target network and edges connecting a target network to the other nine networks together. To further differentiate roles of within- and between-network connectivity in prediction, we assessed the predictive power of between-network connectivity for each network. We constructed a new set of CPMs using functional connections that connect one network with the other nine networks. We performed this analysis iteratively for each of the ten networks. We found that the connectivity of the salience network was again the most predictive of individual attention on average across all prediction cases (*p*s<0.05 FWE corrected using permutation, **Supplementary Figure S10A**). The averaged performance mirrored results from the lesioning analysis; the salience, cerebellum and subcortical networks well predicted individual behaviors (**Supplementary Figure S10B**). Again, to confirm that the lower performance was not driven by the smaller number of between-network connections, we tested the correlation between performance and the number of edges. The results showed no positive correlation between them.

### The salience, frontoparietal and subcortical networks predict universal attention

Given the importance of the salience, subcortical, cerebellum, frontoparietal and visual II networks in predicting individual attentional behaviors, we asked if connectivity between these networks can predict individual behaviors in attention tasks with an accuracy comparable to the original whole-brain models. The CPMs using the connectivity between the five networks fully generalize across different task-related behaviors (**Supplementary Figure S11**). Among the models tested in this analysis, the connectivity of three networks, the salience, frontoparietal, and subcortical networks, were the most important for prediction (**Supplementary Figure S11BC**). Of all models made by selecting any three out of five networks, a model using the salience, frontoparietal, and subcortical networks best predicted individual behaviors on average, and its prediction performance was comparable to the performance of the original whole-brain model. This result suggests that connectivity between the salience, frontoparietal and subcortical networks may be associated with a general component of attention. As a control analysis, we built a model using the other five networks (the medial-frontal, default mode, motor, visual I, and visual association networks) and found that the model’s prediction was less accurate than the previous five-network model or even the three-network (the salience, frontoparietal, and subcortical networks) model (**Supplementary Figure S11B**). This result agrees with the preceding observation that the CPM of a universal attention factor accurately predicts attention task performances (**Figure 4**) and corroborates these networks’ general importance in attention functions.

### C2C modeling successfully generates patterns of attention-related connectomes from resting state fMRI and improves prediction of attentional behaviors

Although the results above demonstrated that task-based connectomes led to better prediction of behaviors compared to rest connectomes, resting scans still have the undisputable advantage of enhancing data retention by reducing the demand on participants, especially in clinical populations. Therefore, we next asked if we could improve individual behavior predictions from rest connectomes by applying connectome-to-connectome (C2C) state transformation modeling (Yoo et al., 2020). We implemented the C2C pipeline to our rest connectomes to generate connectomes of the three attention tasks. The model-generated task connectomes accurately resembled their corresponding empirical task connectomes (**Supplementary Figure S12AB**). The generated task connectomes were more statistically similar to the empirical task connectomes than the empirical rest connectomes were, in terms of the edge-wise strength as well as the spatial pattern of the whole-brain connectome (*p*s=1/1,001). More remarkably, individual attentional behaviors were predicted by the generated task connectomes better than by the empirical rest connectomes alone (*p*s<0.05). The generated connectomes not only captured individual differences (as measured by Pearson’s *r*; **Figure 7B**) but also accurately predicted an individual’s actual behavioral scores (as measured by *q*^*2*^; **Figure 7A**). This result suggests that C2C state transformation is essential to make CPMs generalizable across different cognitive states of fMRI. The results from the movie connectomes are also shown in **Supplementary Figure S12CD and S13**.

**Figure 7.**
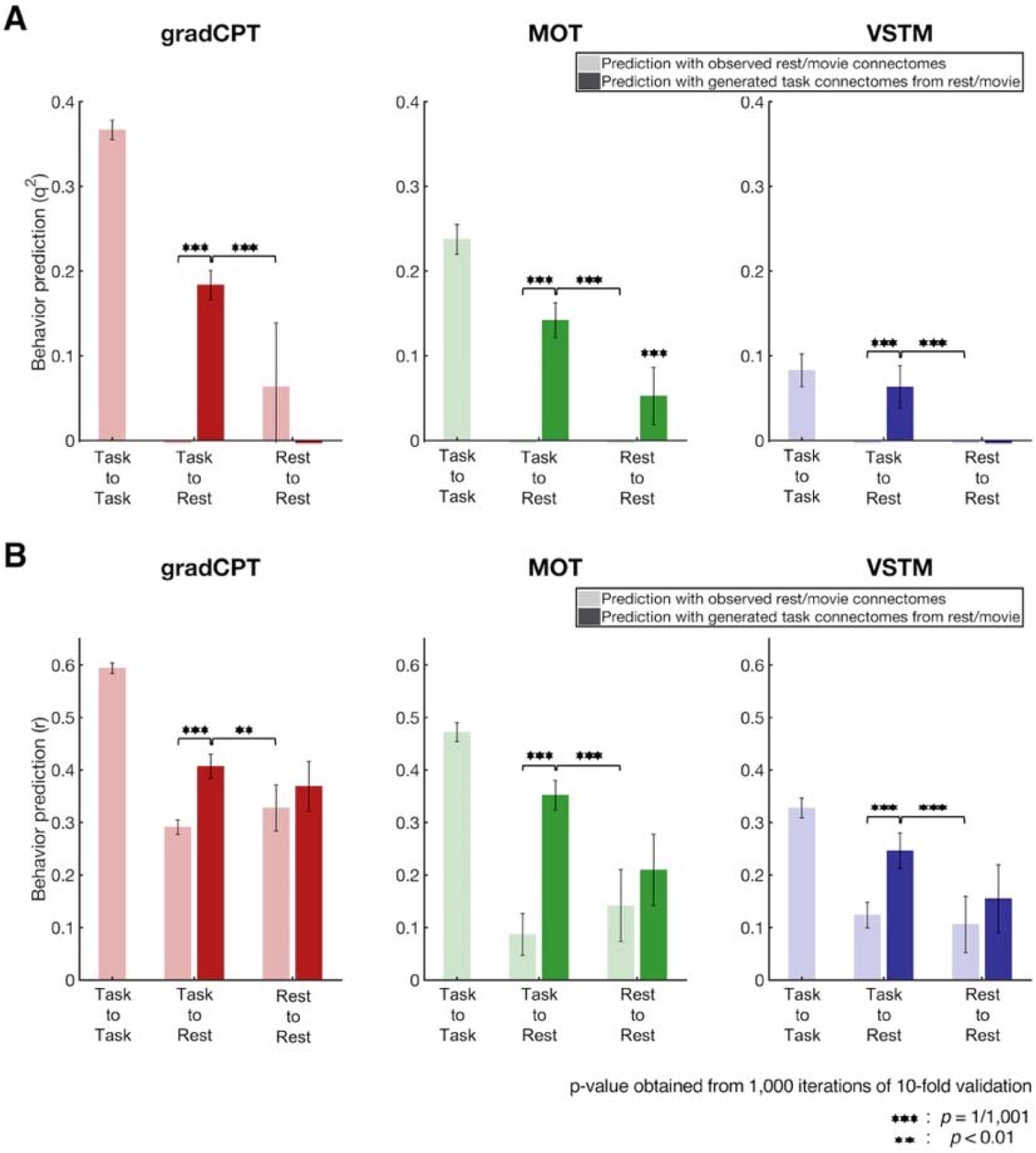
Prediction of individual behaviors with different approaches. **A**. Prediction performance was assessed by prediction q^2^ and negative values were set to zero (i.e., task-to-rest prediction with the empirical rest connectome in all three tasks). Darker bars represent the behavior prediction with task connectomes generated by C2C modeling. The task connectomes generated by C2C modeling from rest data significantly better predicted individual behaviors than empirical rest connectome in all three attention tasks. A darker bar in task-to-rest represents the behavior prediction of a model trained using empirical task connectome and predicted with task connectome generated from empirical rest connectome. A darker bar in rest-to-rest represents the prediction of a model trained using empirical rest connectome and predicted with task connectome generated from rest connectome. **B**. The same result, but prediction performance was assessed by correlation *r*. Error bars represent standard deviation across 1,000 iterations. GradCPT: gradual-onset continuous performance task, MOT: multiple object tracking, and VSTM: visual short-term memory.

### A universal attention model

Unifying the models and findings above, we propose a universal attention model to predict a single attention measure for novel individuals based on resting-state data alone. The universal attention model accurately predicted performance in the three attention tasks based on the rest connectome, and the universal attention model was significantly better than the MOT and VSTM task models applied to rest data with C2C transformations (**Figure 8** and **Supplementary Figure S14A**). When the task-specific models were applied to rest data without C2C transformations, they could not predict individuals’ actual scores at all (q^2^=0 in three task-based models, **Supplementary Figure S14B**), showing the importance of C2C transformation for applying CPMs to rest data.

**Figure 8.**
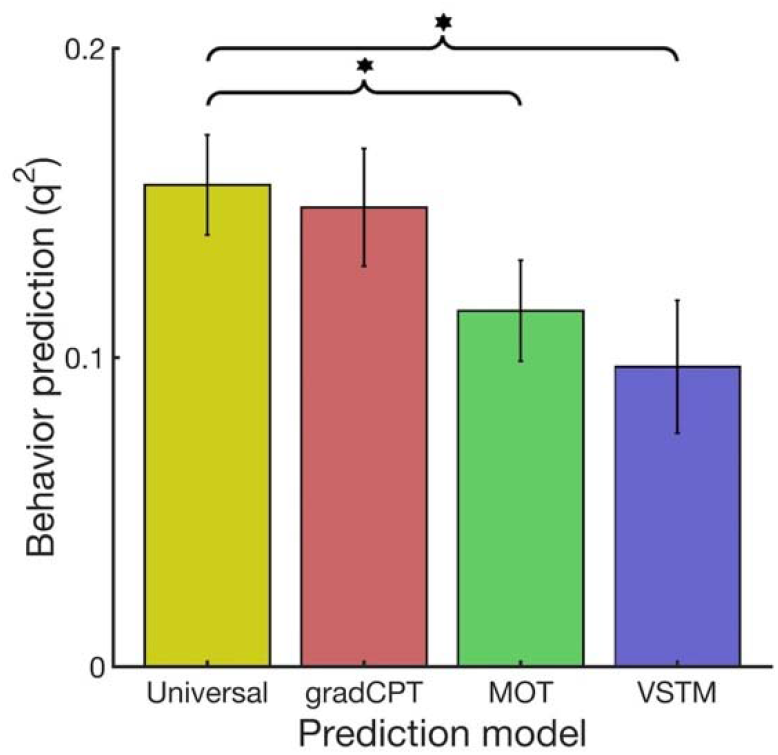
Behavior prediction by the universal attention model. Each task name in *x*-axis represents a single task-based model combined with corresponding C2C model. The universal attention model and three CPMs predict individual behaviors from a resting-state connectome. Behavior prediction represents performance averaged for three tasks prediction. Error bars represent standard deviation across 1,000 iterations. *: The universal model prediction significantly better predicted behaviors than CPMs at p<0.05 from 1,000 iterations. GradCPT: gradual-onset continuous performance task, MOT: multiple object tracking, and VSTM: visual short-term memory.

The universal attention model’s performance was even comparable to task-specific models applied to task data, averaged across three tasks (**Supplementary Figure S15**). Importantly, the universal model generalized across the three tasks better than any of the task-specific models, which made weaker predictions for non-native tasks. The strong predictive power and the higher generalizability of the universal model make it broadly applicable.

### External validation: the universal model generalizes to independent datasets

To further validate the proposed universal attention model’s generalizability and practical applicability, we tested it on three independent datasets. The three datasets comprise rest connectomes and gradCPT scores (*d’*) from 25 adults (Rosenberg et al., 2016a), rest connectomes and ANT scores (*RT variability*) from 41 adults (Rosenberg et al., 2018), and rest connectomes and ADHD Rating Scale-IV scores (DuPaul et al., 1998) from 113 children and adolescents with and without ADHD diagnoses (Rosenberg et al., 2016a) provided by the ADHD-200 consortium (Consortium, 2012).

The universal model successfully generalized to predict attention performance in the three external datasets. The universal model not only captured individual differences in attention function (**Supplementary Figure S17**) but also accurately predicted individuals’ actual scores (**Figure 9**). In contrast, the two CPM models trained using gradCPT fMRI or rest fMRI were much less accurate in predicting actual individual abilities across the different external datasets and measures (**Figure 9**). It should be noted that the gradCPT CPMs were able to detect relative differences in individual attention performance (vs. actual performance) when measured by correlation (**Supplementary Figure S17**). This result demonstrates the practicality of the proposed universal attention model over task-specific CPMs.

**Figure 9.**
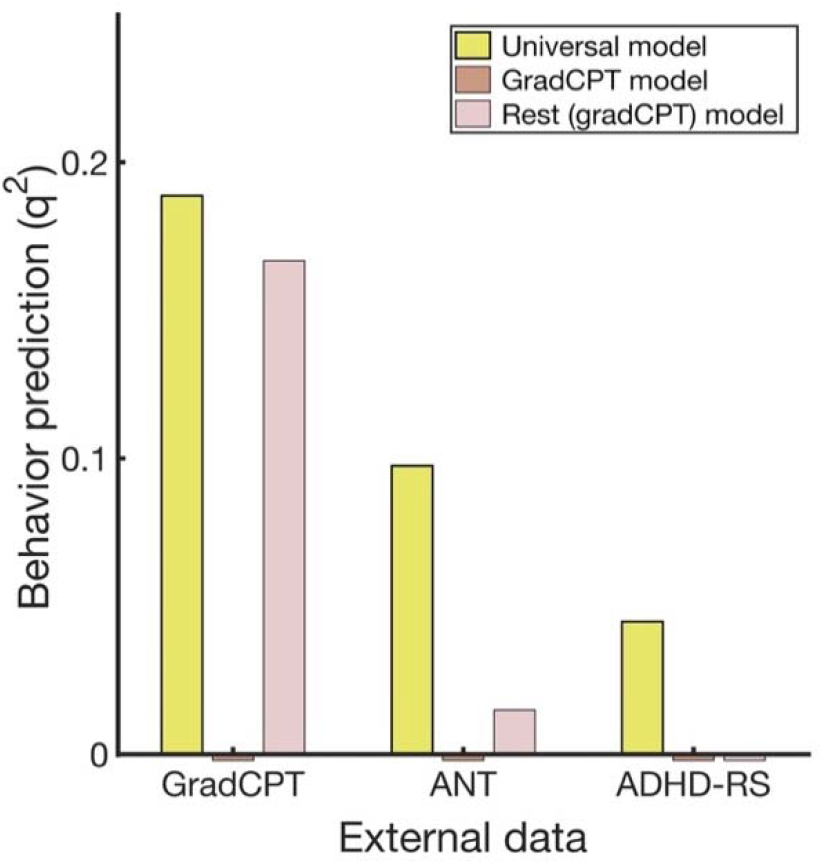
Generalizability of the universal attention model in three independent datasets. **A**. Prediction performance was assessed by prediction q^2^. Negative q^2^ values were set to zero. The universal model (yellow) successfully generalized in different datasets. The universal model accurately predicted individuals’ actual attentional abilities obtained in gradCPT and ANT and assessed by ADHD-RS. In contrast, the CPMs trained using gradCPT fMRI or rest fMRI did not generalize to predict individual abilities that were assessed by different measures in the external datasets.

To confirm that the universal attention model predictions are specific to attention, we performed the following analyses adopted from Rosenberg et al., 2016a, using the ADHD dataset. First, we assessed the correlation-based model performance while controlling for age and IQ measured by the Wechsler Intelligence Scale for Chinese Children-Revised (Dan and Yu, 1990). The universal attention model remained successful, exhibiting a significant positive correlation between observed and predicted scores (*r*=0.223, *p*=1.85×10^−2^), even after controlling for age and IQ. This result indicates that the model indeed measures attention rather than a general cognitive ability or age. Second, to confirm that an individual’s general arousal did not drive the universal model prediction, we assessed the relationship between the model prediction and the hyperactivity-impulsivity. If the proposed universal attention model predicts arousal rather than attention, model-predicted scores should correlate positively with hyperactivity scores that reflect high arousal. Instead, the universal attention model predictions correlated negatively to the hyperactivity scores (*r*=-0.240, *p*=1.03×10^−2^), suggesting that hyperactive individuals have worse attention. Again, this result suggests that the universal attention model captures attentional ability rather than individual arousal.

## Discussion

We developed a suite of whole-brain connectome-based predictive models (CPM) that can predict both a universal (task-general) or multifaceted (task-specific) measure of attention from an individual’s resting-state fMRI data or single-task fMRI data. The network models accurately predicted sustained attention, divided attention and tracking, and working memory capacity. By comparing these models, we uncovered the underlying neural mechanisms supporting a general component across these attentional functions. We observed that patterns of multiple brain networks, including the salience, subcortical, cerebellum and frontoparietal networks, drive accurate prediction of individuals’ attentional abilities across tasks, suggesting that these networks support a universal (general) attention factor. To further enhance the measurement of attention, we applied a novel analysis framework, connectome-to-connectome (C2C) modeling (Yoo et al., 2020), and demonstrated that we can generate the patterns of individuals’ attention task connectomes from their rest connectome alone. More remarkably, the generated task connectomes further improved prediction of individual attention behaviors in either a task-specific or universal manner. The universal attention measure performed better than task-specific models applied to rest data, and it generalized better across the three attention tasks and other external measures, making it a powerful measure with broad utility.

### The generalizability and functional anatomy of attention network models

CPMs successfully generalized to predict performance across three different attention tasks: sustained attention, tracking, and visual working memory (**Figure 2**). For example, a model trained to predict individual behaviors in gradCPT accurately predicted performance in MOT and in VSTM as well. This generalizability of predictive models across different attention tasks suggests shared neural mechanisms of a universal attention factor across the tasks. Previous studies revealed a general attention factor (Huang et al., 2012), and the neural system underlying attention performance in diverse tasks (Wojciulik and Kanwisher, 1999). Going beyond these studies, our CPM approach looks at connectivity patterns that can further predict quantifiable performance across multiple tasks in novel individuals.

The brain networks of the universal attention factor mainly recruited the salience, subcortical, cerebellum, frontoparietal and motor networks (**Figure 4B**). CPMs tuned to the universal attention factor exhibited prediction accuracy and generalizability comparable to the native task models. This was surprising given that the universal attention models utilized significantly fewer connectivity features than the native task models (**Figure 3B** and **Supplementary Table S2**).

To further understand the relative contribution of each canonical brain network for the universal attention factor, we computationally lesioned each network in isolation and examined their impact on whole brain CPM performance. We observed that the salience network, followed by the subcortical and frontoparietal networks, is the most important network in predicting individual attentional behaviors across all tasks in general, that is, exhibiting high generalizability (**Figure 5 & 6 and Supplementary Figure S9 & S10**). The results further show that these networks play a primary role in attention performance across tasks. The involvement of these networks in CPM is in line with previous findings that attention-related tasks induce or modulate a functional engagement of frontal and parietal areas (Corbetta et al., 1995; Hopfinger et al., 2000; Pardo et al., 1991; Sprague and Serences, 2013; Wojciulik and Kanwisher, 1999) and subcortical areas (Coull et al., 2004; Heinze et al., 1994; Wimmer et al., 2015).

Interestingly, we also found that connectivity between the cerebellum and other networks are an informative marker of individual attentional performance (**Figure 5** & **6**). Although the cerebellum is traditionally important for motor control, cerebellar involvement in higher cognition has been proposed over three decades (Gao et al., 1996; Leiner et al., 1986; Petersen et al., 1989; see for reviews, Stoodley, 2012; Strick et al., 2009). Advances in brain imaging techniques started to reveal cerebellar connections with higher association cortices and coactivation with cortical networks in various cognitive tasks (Buckner, 2013). Cerebellar involvement in the specific function of attention has also been well documented; such as, attention-induced cerebellar activation (Allen et al., 1997), attentional modulation on the cerebellar activity (Rees et al., 1997), and attention deficits with cerebellar lesions (Gottwald et al., 2003).

### The validity of the universal attention measure

The universal attention model here was based on three tasks, which cover many fundamental dimensions of attention: sustained attention, divided attention and tracking, and working memory. Amongst the specific tasks, the gradCPT task supports the strongest predictions and generalizability to other tasks here and in other studies (Rosenberg et al., 2016a, 2016b; Yoo et al., 2018). For these reasons, when only one attention task can be conducted in the scanner, we recommend the use of the gradCPT. When only rest data are available, the universal attention model offers the most generalizable measure of attention.

The gradCPT, MOT, and VSTM tasks tested here are only a small sample of the wide variety of attention tasks out there. However, having tested a wider variety of tasks, Huang et al. demonstrated that that there appears to be only one general factor that is shared across them, and we believe that this is what underlies the universal attention measure proposed here (Huang et al., 2012).

As further external validation, our universal attention model generalized to predict attention performance, and not general cognitive ability or arousal, across several novel and independent datasets, multiple measures of attention (gradCPT *d’*, ANT *RT variability*, and ADHD *Rating Scale*), lab or clinical data (ADHD diagnosis), different age groups (adults and children/adolescents), and variable data acquisition sites (two centers in New Haven and one in Beijing) and data processing procedures (AFNI and Bioimage Suite with SPM8). Future work can further validate the universal attention measure across different tasks and datasets such as the Human Connectome Project (Barch et al., 2013), Philadelphia Neurodevelopmental Cohort (Satterthwaite et al., 2014), and Adolescent Brain Cognitive Development Study (Casey et al., 2018).

### Factors affecting prediction accuracy

Our CPMs of different attention functions provided different, at least numerically, prediction accuracy for the three tasks. We attempted to address if any parametric difference in modeling or behavioral measures causes the different accuracy of predictive models. Our results indicate that different reliabilities of the three behavioral measures might have constrained the prediction accuracy of models. Individual behaviors in gradCPT were most reliable, and the model prediction of gradCPT performance was the most accurate (**Supplementary Figure S1**). Individual behaviors in VSTM were least reliable, and the model prediction of VSTM performance was the least accurate, although it was still significant. Other factors – the degree to which the general factor explains individual behaviors in each task (**Supplementary Table S4**), the number of predictive features, the number of available subjects, and scan duration – were uncorrelated with prediction accuracy (or controlled to be consistent across tasks in experimental design). This result underlines the obvious importance of measuring behaviors accurately during fMRI scanning.

### Functional connectome during the movie-watching and the resting states

The idea of using naturalistic stimuli during fMRI acquisition has attracted research interest. Movie-watching paradigms of fMRI were demonstrated to amplify individual differences of brain organization (Vanderwal et al., 2017) and to better capture individual differences in intelligence than predictions based on resting state fMRI (Finn and Bandettini, 2020). Expanding on these reports, we demonstrated improved predictions of individual attention performance from movie-watching scans. However, it remains unclear why movie-watching data supports better predictions than rest data. One possibility is that movie watching induces attentional engagement of participants, making their functional connectomes more similar to attention task connectomes. To address if this was the case, we estimated the spatial similarity of movie-watching connectomes to attention task connectomes and compared it to the similarity of resting-state connectomes to attention task connectomes. We observed that these similarities were not significantly different, suggesting that being engaged in a movie does not simply make individual connectomes more similar to attention-task connectomes. Another possibility is that movie watching aligns individual brain organization to be more consistent with each other (responding to the same visual stimulus), thus reducing individual variability from random mind wandering, which should be widely different across subjects and between sessions. Previous studies reported that movie-watching paradigms improved reliability and individual identification (Meer et al., 2020; Wang et al., 2017). To examine this in our dataset, we estimated and compared intra- and inter-individual similarity of connectomes at resting-state and at movie-watching. fMRI time-volumes made available to the connectome construction was matched between two states: 300 TRs from each session. We observed, however, that the connectome similarity (both within and between individuals) during movie-watching was only numerically higher than the resting-state and we did not see any statistical difference. This suggests that subjects may exhibit a substantial difference in processing the same stimuli based on own experiences and knowledges, and the difference might be reflected in across the whole brain.

### Practical utility and potential applications

To enhance the practicality of the current approach and models, we extended CPM modeling with a novel method called connectome-to-connectome (C2C) modeling. We showed that C2C modeling accurately predicted the individual attention-task connectomes from their rest connectomes, and notably, it enhanced behavioral predictions better than using the rest connectome alone.

The C2C framework should have significant practical and clinical promise. Compared to task scans, rest scans are easier to collect consistently across studies and sites. For example, clinical populations may have difficulty performing certain tasks (Pujol et al., 1998). Instead, researchers or clinicians can obtain rest scans from patients because of its simplicity and minimal demands (Bullmore, 2012). This is one of the main reasons why resting-state fMRI has gained much popularity in clinical and other neuroimaging studies. However, relative to task data, rest data have reduced accuracy in characterizing individual traits and behaviors (Greene et al., 2018; Jiang et al., 2020; Tomasi and Volkow, 2020; Yoo et al., 2018), perhaps due to unconstrained mind-wandering that may result in more variable mental states from scan to scan and from subject to subject. The C2C modeling improves the diagnostic value of rest scans, easing the burden of conducting multiple scans or trying to standardize tasks across individuals.

In this study, we demonstrated the applicability of the C2C framework in studying attention functioning. Our approach of combining CPM and C2C modeling should be useful in studying other mental abilities such as memory and fluid intelligence, and related neuropsychiatric disorders such as attention-deficit/hyperactivity disorder and dementia. Our approach can derive estimates of multiple cognitive measures from a single rest scan, analogous to how physicians can assay multiple health measures from a single blood sample. The universal attention measure is proposed as a standard to quantify and compare attention functioning across individuals and testing sites.

## Conclusion

We developed a suite of connectome-based models that can predict a profile of individual attentional abilities from a single fMRI scan. The task fMRI-based models accurately predicted attentional components of sustained attention, divided attention and tracking, and working memory capacity. We observed that patterns of multiple brain networks, including the frontoparietal, salience, and subcortical networks, drive the accurate prediction of individuals’ attentional abilities across tasks, suggesting that these networks underlie a general factor of attention. The combined approach of CPM and C2C allows these complex profiles of an individual’s cognitive and behavioral abilities to be estimated from commonly collected rest scans.

## Acknowledgement

This project was supported by National Institutes of Health grant MH108591 and by National Science Foundation grant BCS1558497.

## Methods

### Subjects and experimental designs

One hundred and twenty-seven right-handed, neurologically healthy individuals with normal or corrected-to-normal vision participated in a two-session fMRI study (80 female, ages 18 to 35 years, mean=23.15 years, SD=4.43). Data from thirty-three participants were excluded from the analysis due to excessive head motion (>3 mm maximum head displacement and >0.15 mm mean framewise displacement [FD]) during fMRI scanning, task performances with lower or higher than 2.5 standard deviations from the group mean in both sessions, or low imaging data quality. The remaining 94 individuals with all behavioral and imaging data were included in the main analysis (61 female, ages 18 to 35 years, mean=23.07 years, SD=4.51). Two fMRI sessions were separated by approximately two weeks (mean=17.27 days, SD=20.29).

Each fMRI session started with an anatomical magnetization prepared rapid gradient echo (MPRAGE) followed by 10-min resting-state runs (two runs in session 1 and one run in session 2) and a 7:16-min watching-movie run (*Inscapes*, Vanderwal et al., 2015). Afterwards, all participants performed three 10-min attention-related tasks while they were in the scanner. The three tasks were the gradual-onset continuous performance task (gradCPT), multiple object tracking (MOT), and visual short-term memory task (VSTM). The order of these tasks was counterbalanced across participants and sessions. An additional task, either the Attention Network Task (ANT; Fan et al., 2005) or an *n*-back task, was collected after completing the three main tasks in session 2, but these tasks are not included in this study. All participants provided written informed consent approved by the Yale University IRB and were paid for their participation.

### Three attention tasks

Participants performed three attention-related tasks in the scanner. Task performance was assessed with sensitivity (*d’*), accuracy (%), and working memory capacity (Pashler’s *K)* for gradCPT, MOT, and VSTM, respectively. To calculate task performance scores, we averaged scores from the two sessions for each task. For those who had only one session that met our data inclusion criteria (29 subjects for gradCPT, 25 subjects for MOT, and 22 subjects for VSTM), we used the task score of the available session in the analysis.

#### Gradual-onset continuous performance task

The gradCPT is a task that measures sustained attention and inhibitory control (Esterman et al., 2013; Rosenberg et al., 2013). In this 10-min task, participants saw grayscale photographs of scenes gradually transitioned from one to the next. The scenes consist of city scenes that appear in 90% of the total trials and mountain scenes that appears in only 10% of the total trials. Each scene transitioned every 800ms, and participants were asked to respond every time they saw a city scene by pressing a button with their right index finger and withhold responses to the mountain scenes. The task consisted of 740 trials (.8 s each). Sensitivity (*d*’) was calculated to assess task performance.

#### Multiple object tracking

MOT measures divided attention, tracking, working memory capacity, spatial attention, inhibition, and sustained selective attention (Pylyshyn and Storm, 1988). In this 10-min task, participants tracked multiple target objects while all stimuli were moving. At the beginning of each trial, participants were presented with 12 randomly spread identical white discs on the screen. For each trial, three or five discs among the 12 flashed green and turned back to white, designating them as the target discs of that trial, while the remaining non-target discs remained white. All of the 12 discs then moved around the screen for 5,000 ms, and then one of the 12 discs was probed. Participants were instructed to press a button with their right index finger if the probed disc was one of the original targets and press with their right middle finger if it was not. The task consisted of 56 trials (10.3 s each), and performance was assessed by a percent accuracy.

#### Visual short-term memory task

VSTM measures visual working memory capacity that stores visual information (Luck and Vogel, 1997). In this 10-min task, participants saw discs of same size but different colors on the screen for 100ms and were asked to remember the colors of individual discs. The number of discs for each trial varied from two to eight (two, three, four, six, or eight discs). The stimuli were replaced by a fixation mark for 900ms, and the discs reappeared with or without color changes. Participants were instructed to press a button with their right index finger if they detect any color changes between two appearances of the discs and press the other button with their right middle finger if no change had occurred. Participants had 2,000ms to respond. The task consisted of 160 trials (3.6 s each). For half of the total trials, original discs were replaced by different colors of discs, and for the other half of the trials, the original discs remained unchanged. Visual working memory capacity was assessed with Pashler’s *K* (Pashler, 1988).

### Behavioral analysis: Individual performance in three attentional tasks

Given that all tasks required attentional ability, individuals who performed well in one task were expected to perform well on others. In order to confirm how behaviors in different attentional tasks are related, we computed Pearson’s correlation between individual performance scores on every pair of tasks, resulting in three between-task similarity metrics: between gradCPT and MOT, gradCPT and VSTM, and MOT and VSTM.

The between-task similarity estimate should be constrained by the reliability of the behavioral measures we adopted. Therefore, prior to computing a similarity of individual performance on the three tasks, we assessed the reliability of each behavioral measure. For this analysis, we only used subjects who had acceptable behavioral scores from both sessions for each task, resulting in a different number of available subjects for each task; 65 subjects for gradCPT, 69 subjects for MOT, and 72 subjects for VSTM. Within these subsamples, we estimated Pearson’s correlation of individual performance between the two sessions for each task.

### MR imaging parameters and preprocessing

MRI data were collected at the Yale Magnetic Resonance Research Center and the Brain Imaging Center at Yale with 3T Siemens Prisma system and 64-channel head coil. A high-resolution MPRAGE was collected at the beginning of each session with the following parameters: TR = 1800 ms, TE = 2.26 ms, flip angle = 8°, acquisition matrix = 256 × 256, in-plane resolution = 1.0 mm^2^, slice thickness = 1.0 mm, 208 sagittal slices. 10-min resting-state scans, two scans in session 1 and one in session 2, were collected after MPRAGE followed by an Inscape movie-watching run (7:16 min). After these passive viewing scans, participants performed three 10-min main attention-related tasks, gradCPT, VSTM, and MOT, with a button box in their right hand. Each of the three tasks and resting-state scans included 600 whole-brain volumes acquired using an EPI sequence with the following parameters: TR = 1,000 ms, TE = 30 ms, flip angle = 62°, acquisition matrix = 84 × 84, in-plane resolution = 2.5 mm^2^, 52 axial-oblique slices parallel to the AC-PC line, slice thickness = 2.5 mm, multiband 4, acceleration factor = 1. This information was also provided in Rosenberg et al. (2020), which analyzed a subset of the current dataset; 49 subjects with two usable gradCPT runs at the time of the study (Rosenberg et al., 2020)

Collected data were preprocessed with AFNI (Cox, 1996). The preprocessing procedure included the following steps: Removing the first three volumes; censoring of volumes containing outliers in more than 10% of voxels; censoring of volumes for which the Euclidean norm of the head motion parameter derivatives are greater than 0.2 mm; despiking; slice-time correction; motion correction; regression of mean signal from the CSF, white matter, and whole brain and 24 motion parameters. FMRI data were aligned to the high-resolution anatomical image (MPRAGE) and normalized to MNI space. All the following main analyses were performed in MATLAB R2016b.

### Whole-brain connectivity matrix

Functional network nodes were defined using a 268-node whole-brain functional atlas, which covers the cortex, subcortex, and the cerebellum (Shen et al., 2013). We excluded 22 nodes (due to imperfect acquisition of fMRI data on these areas from at least one subject), resulting in 246 nodes analyzed in this study. For each participant, an averaged time-series signal was calculated for each node, and Pearson’s correlation between all possible pairs of the 246 nodes were computed. The pairwise correlations were then Fisher z-transformed, resulting in a 246×246 symmetrical whole-brain functional connectivity matrix (30,135 unique edges). We calculated the connectivity matrix for each session separately and averaged them across two sessions for the final analysis. For those who had only one session that met our data inclusion criteria (29 subjects for gradCPT, 25 subjects for MOT, and 22 subjects for VSTM), we used the connectivity matrix from the available session in the analysis. Every individual had five connectivity matrices including three attention-related, one resting-state, and one movie-watching.

### Brain-based prediction of individual behaviors across three attention tasks

Previous studies demonstrated that CPMs accurately predict individual gradCPT performance from both task-related and resting-state fMRI connectivity (Rosenberg et al., 2016a; Yoo et al., 2018). Extending these previous studies, we asked in the current study whether different attention tasks lead to different predictive models that are generalizable across multiple attentional behaviors. To this end, we constructed nine CPMs using different combinations of imaging and behavior data; that is, three different cognitive states of fMRI scans (performing one of three attention tasks, resting-state, or movie-watching) and three different attentional behaviors (gradCPT, MOT, or VSTM). A detailed procedure of CPM modeling and generalizability tests are described in the following sections.

#### Connectome-based predictive modeling (CPM)

We constructed and validated CPMs using a leave-one-out cross-validation (LOOCV). In building CPMs, we held one subject out for model testing, with 93 participants in the training set. In training the CPM model, we first selected features (edges) that were significantly correlated with individual behaviors in a target task (Pearson’s, *p<0*.*05*). These features yielded both positive and negative edge masks depending on the signs of their correlation with behavior. For each subject in the training set, two networks’ strengths (one from the positive and the other from the negative network) were measured by averaging their respective connectivity strengths. Then, we fitted a general linear model between task performance (a dependent variable) and the two network strengths (independent variables). Once the two network masks and a general linear model were constructed, we applied the CPM to the held-out testing subject. The CPM estimated two network strengths for the test subject and predicted the subjects’ task performance from their network strength measures. Every subject was iteratively used as a test sample in LOOCV.

We previously demonstrated that CPMs are robust against the choice of feature selection threshold within the range of traditional statistical significance (e.g., *p* of 0.05 ∼ 0.001) (Yoo et al., 2018, 2019). We tested a similar range of selection thresholds in the current study and confirmed that the results remained similar across the range.

#### CPMs of different attentional tasks

Here, we tested whether we could construct successful CPMs for each attention task, and whether CPMs of attentional tasks generalized to predict individual behaviors across different tasks. To address these questions, we constructed a total of nine CPMs. These models were constructed using different combinations of three cognitive states of fMRI scans (task-performing, resting-state, or movie-watching) and three attentional behaviors of target tasks (gradCPT, MOT, or VSTM) (**Table 1**). The nine CPMs include seven new models and two models (model 1 and 4 in **Table 1**) that were introduced in our previous studies (Rosenberg et al., 2016a; Yoo et al., 2018).

We assessed the performance of these nine models in predicting individual behaviors of interest. First, we tested model performance of a *within-task prediction*, for example, training the model using VSTM fMRI and VSTM behavioral task scores and testing it to predict the VSTM behavior from VSTM fMRI of the held-out subject. To examine model generalizability across different tasks, we then assessed prediction performance across tasks and fMRI, in a *cross-task prediction analysis*. For example, the model 1 in **Table 1** was trained using gradCPT fMRI and gradCPT behavior. We applied this model to a testing subject’s MOT fMRI to predict the MOT score of the subject to assess the generalizability of the model to MOT fMRI and MOT behavior. In this way, we tested each model in each of the nine fMRI–behavior pairs, yielding 72 cross-task predictions along with nine within-task predictions from nine predictive models (as shown in **Figure 2**, diagonal elements correspond to *within-task predictions* and off-diagonal correspond to *cross-task predictions*).

We assessed each model’s prediction performance by correlating model-predicted individual task scores and observed task scores. A significant positive correlation indicates that the model successfully predicts individual differences in behavioral performance. We also estimated prediction q^2^, based on normalized MSE, to further validate model prediction (Scheinost et al., 2019). The prediction q^2^ represents a model’s numerical accuracy in predicting an individual’s actual behavioral score compared to simply guessing their mean behavior. The prediction q^2^ serves as a complementary measure of the correlation-based performance measure that describes the accuracy in predicting individual differences between behavioral scores.

Behavioral scores were z-scored for each task before predictive modeling. Z-scoring was essential to compare predictions across multiple tasks with incompatible scoring scales. We used raw scores only in the visualizations of within-task prediction results with scatter plots.

#### Significance testing of CPMs’ behavioral prediction accuracy

We evaluated the significance of model performance using permutations. We ran 1,000 permutations to construct 1,000 null models for the 81 model predictions. In each permutation, individual performances were randomly shuffled, and the null CPMs were trained and tested with connectivity matrices and the shuffled performances for 81 prediction cases. We assessed the performance of the null models by correlating the model-predicted behavioral scores and the shuffled scores. The same evaluation was applied to the real model, except shuffling individual behaviors.

Importantly, we used permutations to correct for multiple tests. To do this, we first divided 81 predictions into nine divisions based on cognitive states of fMRI of training and testing datasets (1: task fMRI to task fMRI prediction; 2: task to rest; 3: task to movie; 4: rest to task; 5: rest to rest; 6: rest to movie; 7: movie to task; 8: movie to rest; 9: movie to movie). Then, we further subdivided each division into two groups, one group of three within-task prediction cases and the other of six cross-task prediction cases, resulting in 18 case groups in total. We corrected for multiple comparisons for each case group separately. In each permutation run, the maximum null performance was selected for each case group, to yield 1,000 maximum performance null models for each group. We compared observed model performance with the 1,000 max null model of the corresponding group. The FWE-corrected significance of the observed model performance was calculated as *p* = (1 + the number of the null performances better than the observed model performance) / 1,001.

### Generalizing CPMs while controlling for behavioral correlations between tasks

In the previous analysis, we examined the generalizability of CPMs across different tasks. Successful generalization, however, may be considered trivial, given the significant correlation of individual performances between tasks. To address this issue, we controlled for behavioral correlations across tasks in assessing model performance. In the original model building, feature edges were selected based on the significance of their association with a behavior of the target. We constructed additional models while controlling for the correlated behaviors. In these models, among the selected features in the original model, we excluded edges that were significantly correlated with behaviors in either of the other two tasks. In other words, these models only included predictive edges that were specific to one task. We assessed the performance of these models as described in *<u>Significance testing of CPMs’ behavioral prediction accuracy</u>*.

We further examined the generalizability and specificity of the original models by applying partial correlations. In this case, we re-assessed the original models’ prediction accuracy using partial correlations between model-predicted and observed behavioral scores while controlling for scores in non-target tasks. We assessed the performance of the models as described in *<u>Significance testing of CPMs’ behavioral prediction accuracy</u>*.

### Predictive anatomy of attention CPMs

We explored the predictive anatomy of CPMs to reveal the anatomical basis of the attention tasks. Each iteration of LOOCV provided positive and negative network masks for each model. We extracted the most robust edges that appeared in every LOOCV iteration of each modeling. These robust predictive edges were then visualized with ten canonical networks (medial frontal, frontoparietal, default mode, motor, visual I, visual II, visual associations, salience, subcortical, and cerebellum; Finn et al., 2015; Noble et al., 2017).

### Controlling for confounding factors

#### The number of predictive features in CPM modes

Each attention task CPM has a different number of predictive edges, which may induce differences in predictive power. To address this concern, we controlled the number of features, revised the feature selection threshold, and constructed three new task-based CPMs. In the original modeling, we selected all edges that were significantly associated with a behavior of interest. In the current analysis, we selected the top 1,000, 5,000, and 10,000 edges that were most significantly associated with a behavior as features. By doing this, all models contained the same number of feature edges. We tested the performance of the models in the same way as the original modeling, including significance testing with FWE correction.

#### Effect of head motion on behavior prediction

As described earlier, we took into account head motion in this study 1) by censoring high motion fMRI frames in each data (described in ***MR imaging parameters and preprocessing***), 2) by excluding subjects with excessive head motion (described in ***Subjects and experimental designs***) and 3) by regressing out head motion parameters during MR preprocessing (described in ***MR imaging parameters and preprocessing***). These procedures, however, might not fully eliminate the influence of head motion on the estimations of associations between behavior and brain connectivity, thereby affecting connectome-based predictions of behavior. To address this issue, we re-assessed the prediction accuracy of nine original CPMs, taking head motion into account. We ran partial correlations between model-predicted and observed behavioral scores while controlling for head motion from the scan of the target task. We confirmed that the maximum FD values of all five fMRI data (three tasks, one rest, and one movie) did not correlate with behavioral performance in any of three tasks. Hence, we controlled for mean FD as a head motion estimate of task scans.

### CPMs of a general attention component to develop a universal measure

To examine how well a general component of attention can explain behaviors on a variety of attentional tasks, we built CPMs using only functional connections that were associated with all three task behaviors. In the original task-specific models, feature edges were selected based on the significance of their association with a behavior of the target task. In this variation, among the selected features in the original models, we further selected edges that were significantly correlated with behaviors in the other two tasks (the intersection of edges). By doing so, we constructed one set of network masks of which respective edge strengths were correlated with all three task behaviors across individuals. Hence, the number of connectivity features was significantly fewer than in the original models. We trained and validated these “intersection” models using task fMRI in the same way as the original models, except for this feature selection step.

To reveal a set of connectivity features that supports the general factor of attention across all three fMRI tasks, we tracked common predictive connectivity at the network level. We extracted the predictive anatomy of three task-based CPMs of the universal attention factor (refer to ***Predictive anatomy of attention CPMs*** for details). Since each model had a small number of connectivity features, we loosened the threshold, selecting robust edges that appear in at least 75% of LOOCV iterations (i.e., 750 iterations out of 1,000). We then tracked predictive connectivity features that the three task-based models shared in common.

In addition, we trained nine additional predictive models to utilize a set of connections correlated with any of three task behaviors as features (all task-related edges). Hence, these “union” models have larger numbers of predictive feature edges compared to the original models. We validated these models in the same ways as above, except for the feature selection step.

### Role of canonical brain networks in attentional behaviors

#### CPMs with computational lesion in brain networks

Next, we investigated which brain network is the most predictive of all three attention task scores. To assess the network-wise importance in behavioral prediction, we divided 246 brain nodes into ten canonical networks (the medial frontal, frontoparietal, default mode, motor, visual I, visual II, visual associations, salience, subcortical, and cerebellum) and computationally lesioned all nodes of a given network. Then we constructed and evaluated the nine CPMs of the reduced size of the connectivity matrix after the lesioning. We repeated this procedure by lesioning each network iteratively. We assessed the performance of the models in the same way as the original model assessment, including significance testing with FWE correction. We restricted this analysis to task-related connectivity matrices which yielded to provide a successful prediction in the preceding analysis.

#### CPMs using connectivity of brain networks: within-network connectivity and between-network connectivity

In addition to the computational lesioning, we performed complementary analyses to examine the predictive power of brain networks. In this analysis, we restricted CPMs to use functional connections of only one brain network instead of the whole-brain connectivity. This analysis was further separated into two parts. First, we constructed CPMs based on connectivity within each brain network. Second, we constructed CPMs based on connectivity of one target network to the other nine networks. Hence, the first analysis was to examine predictiveness of within-network connectivity, and the second analysis was to examine predictiveness of between-network connectivity. We assessed the performance of the models in the same way as the original modeling, including significance testing with FWE correction.

### Generating attention connectomes from resting-state data using connectome-to-connectome state transformation modeling

We utilized a novel method from our lab called connectome-to-connectome state transformation modeling (C2C modeling; Yoo et al., 2020) to facilitate the estimation of attention-task connectomes and to improve behavioral predictions from resting-state data alone. In the previous analyses, we examined the generalizability of CPMs across multiple attention tasks. However, predictions of individual scores are typically impaired when the testing sample of fMRI data is different from the training samples. The C2C framework estimates task connectomes from the rest connectome or movie-watching connectome, and by employing the C2C approach to generate three attention task connectomes from rest data, we can improve individual behavioral predictions.

The C2C model works in three steps. First, the model extracts subsystems from the whole-brain resting-state connectome of individuals. The model, then, transforms the extracted subsystems to estimate task-specific subsystems. Finally, the model constructs whole-brain task-specific connectomes from estimated subsystems. More specifically, the C2C model first defines and extracts state-specific subsystems and their scores separately for the resting-state and task-related state using two principal component analyses (PCA). We applied one PCA on the rest connectomes of individuals in the training set. This corresponds to the first step of the C2C model described above. We applied another PCA separately on these same individuals’ gradCPT connectomes. This second PCA provides a reconstruction the whole-brain task connectome from the generated task subsystems, corresponding to the third step of the C2C model. Then, we employed partial least square regression to estimate the transformation of subsystems from the resting-state to the gradCPT state. The PCA-extracted subsystem scores of the resting and gradCPT states were put into the regression. This corresponds to the second step.

In this analysis, we constructed three C2C models to predict whole-brain connectomes of the three attention tasks from the rest connectomes. We assessed the success of task connectome generation in 10-fold cross-validation. We held out one fold (9 or 10 subjects) for model validation and used nine folds to train C2C models. For model validation, we calculated the similarity between the model-generated connectomes and observed task connectomes and the similarity between observed rest connectomes and observed task connectomes, using spatial correlation. We also estimated the root mean square difference between the model-generated and observed task connectomes and compared it with a difference between the observed rest and task connectomes. Finally, we used the model-generated task connectomes to predict individual behaviors in the three attention tasks. We compared prediction accuracy of the generated task connectomes with the accuracy of the observed rest connectomes. Here, CPMs and C2C models were trained in the same training partition of 10-folds and tested in the held-out fold. That is, all naive CPMs to which we compared C2C-combined models were re-constructed and validated in 10-fold cross-validation with 1,000 iterations. We confirmed identical CPM performances between LOOCV and 10-fold cross-validation. We ran the same C2C modeling and comparison procedure using movie data.

### Building a universal attention model

To maximize the practical utility of our suite of attention prediction models described above, we developed a universal attention model that integrates the multiple CPM and C2C models to (1) generate a universal attention connectome from a rest connectome and (2) predict universal (general) attention performance in novel individuals.

The first step for model training was to generate for each participant a universal attention connectome that combines the edges from the individual’s three attention task connectomes. There are several methods for doing so, described at the end of this section, and we chose the method that selects the edge with the highest absolute strength across the three tasks. To do this consistently across individuals, we first computed a group-average attention task connectome by averaging the edge strength from all the training participants for each edge in each task connectome. Then to select which task edge to use for the universal attention connectome, we compared the absolute mean strength for each edge across the three average task connectomes. For example, if the average gradCPT connectome showed the maximum absolute strength for a particular edge, relative to the absolute edge strength in the other average task connectomes (MOT and VSTM), then we assigned the gradCPT edge strength to be the representative edge in the universal attention connectome lookup table (**Supplementary Figure S16**), which specifies which of the task edges to use. Then for each participant, we used this population-level the universal attention connectome lookup table to generate the individual’s universal attention connectome as a mosaic of the empirical edge values pulled from the individual’s three task connectomes.

The resulting individual universal attention connectome can then be fed into the CPM and C2C pipeline like any other individual connectome. We trained one CPM to predict the averaged task score from the representative connectome: the average score across the three tasks is significantly correlated with their first principal component (*r*=0.9998, **Supplementary Table S4**), suggesting that the average performance reflects a universal (general) attention factor (for the three tasks). This CPM selected all the connectivity edges that correlated with any of three task scores as predictive features. We also trained one C2C model that can estimate individual universal attention connectome from novel rest connectomes. Once trained, the universal attention model, combining the CPM and C2C models, can predict a novel participant’s universal (general) attention performance from a single rest connectome. This model was constructed and validated using 10-fold cross-validation.

We explored different variants of ways to build the universal attention model, and the primary model described above was chosen based on simplicity and performance, although performance did not vary significantly between models. To combine the task connectomes into the universal attention connectome, one could average the edges, concatenate the three task connectomes, choose task edges that showed the largest variance across participants, or choose task edges that showed the maximum absolute strength for each participant (without consistency across them). These different methods showed only small numerical differences in performance with the averaging method showing the lowest performance. For feature selection, we also considered edges that correlated with all three task scores (the intersection of features across tasks), rather than with any of the three task scores (the union of features across tasks). The intersection performed slightly worse, so we chose the union model. Finally, we tried predicting behavior using three inner linear regressions to predict the three task scores and then average them for the universal measure; this method performed similarly to the primary model. We tested all the different combinations of these modeling choices and settled on the primary model described above because of its simplicity over the other models, again noting that prediction performance of the different model parameter choices was similar.

### External validation

Lastly, we validated the proposed universal attention model in three independent external validation datasets, two locally obtained (our lab) and one publicly available, to assess the attention models’ generalizability and practical applicability. One local dataset included rest fMRI connectomes and gradCPT performance (sensitivity, or *d’*) from 25 young adults (Rosenberg et al., 2016a), and the other local set included rest fMRI connectomes and ANT performance (correct-trial response time [RT] variability) from 41 young adults (Rosenberg et al., 2018). Task connectomes from these two studies were not analyzed here. Imaging data from these two datasets were acquired using similar parameters to the current study but pre-processed differently. Detailed descriptions of task paradigm, imaging parameters and preprocessing, and data inclusion criteria can be found in Rosenberg et al., 2016a (gradCPT) and Rosenberg et al., 2018 (ANT). All participants in the two datasets gave written informed consent in accordance with the Yale University Human Subjects Committee and were paid for their participation. The open dataset included rest fMRI connectome and ADHD Rating Scale-IV (DuPaul et al., 1998) from 113 children and adolescents with and without ADHD diagnoses, including 75 typically developing controls, from the Peking University site (Rosenberg et al., 2016a) provided by the ADHD-200 consortium (Consortium, 2012). Detailed descriptions of imaging parameters and preprocessing, and data inclusion criteria can be found in Rosenberg et al., 2016a and online at the International Neuroimaging Data-Sharing Initiative (http://fcon_1000.projects.nitrc.org/indi/adhd200/). Each participant’s parent provided informed consent, and all children agreed to participate in the study. The data collection was approved by the Research Ethics Review Board of Institute of Mental Health, Peking University. The variable data collection and analysis procedures across multiple studies and sites enabled a strong test of our model’s generalizability. That is, if our universal attention model successfully predicts attentional behavior across the diverse datasets, then it emphasizes our model’s practical utility.

For external validation analyses, we restricted our predictive network to 230 nodes by removing nodes missing in the ADHD data, resulting in 26,335 (=230×229/2) unique edges in each connectome. Otherwise, the gradCPT task-based CPM model, the rest CPM trained to predict gradCPT score, and the universal attention model were identical to the models tested throughout the present study (n=94). We tested these three models in the three independent external datasets where the attention measures were *d’* in gradCPT, *RT variability* in ANT, and *ADHD Rating Scale* in ADHD. The attention measures were z-scored in each dataset. We reversed the sign of z-scores of the ANT RT variability scores and ADHD Rating Scale-IV scores so that a higher score represents better attention performance. Model performance was assessed by the prediction q^2^ and correlation.

## Data and Code availability

Scripts for the predictive model construction are available for download at https://github.com/rayksyoo/UniversalAttention. Scripts for the other (statistical) analyses are available from the corresponding author upon request. Raw task and rest fMRI data used in the primary analyses will be made available at NIMH Data Archive.

## References

Allen, G., Buxton, R.B., Wong, E.C., and Courchesne, E. (1997). Attentional activation of the cerebellum independent of motor involvement. Science (80-.). 275, 1940–1943.

Avery, E.W., Yoo, K., Rosenberg, M.D., Greene, A.S., Gao, S., Na, D.L., Scheinost, D., Constable, T.R., and Chun, M.M. (2019). Distributed patterns of functional connectivity predict working memory performance in novel healthy and memory-impaired individuals. J. Cogn. Neurosci. 32, 241–255.

Barch, D.M., Burgess, G.C., Harms, M.P., Petersen, S.E., Schlaggar, B.L., Corbetta, M., Glasser, M.F., Curtiss, S., Dixit, S., Feldt, C., et al. (2013). NeuroImage Function in the human connectomelJ: Task-fMRI and individual differences in behavior. Neuroimage 80, 169–189.

Beaty, R.E., Kenett, Y.N., Christensen, A.P., Rosenberg, M.D., Benedek, M., Chen, Q., Fink, A., Qiu, J., Kwapil, T.R., Kane, M.J., et al. (2018). Robust prediction of individual creative ability from brain functional connectivity. Proc. Natl. Acad. Sci. U. S. A. 115, 1087–1092.

Biederman, J., Newcorn, J., and Sprich, S. (1991). Comorbidity of attention deficit hyperactivity disorder with conduct, depressive, anxiety, and other disorders. Am. J. Psychiatry 148, 564–577.

Buckner, R.L. (2013). The cerebellum and cognitive function: 25 years of insight from anatomy and neuroimaging. Neuron 80, 807–815.

Bullmore, E. (2012). The future of functional MRI in clinical medicine. Neuroimage 62, 1267– 1271.

Cai, H., Chen, J., Liu, S., Zhu, J., and Yu, Y. (2020). Brain functional connectome-based prediction of individual decision impulsivity. Cortex 125, 288–298.

Casey, B.J., Cannonier, T., Conley, M.I., Cohen, A.O., Barch, D.M., Heitzeg, M.M., Soules, M.E., Teslovich, T., Dellarco, D. V., Garavan, H., et al. (2018). The Adolescent Brain Cognitive Development (ABCD) study: Imaging acquisition across 21 sites. Dev. Cogn. Neurosci. 32, 43– 54.

Chun, M.M., Golomb, J.D., and Turk-Browne, N.B. (2011). A Taxonomy of External and Internal Attention. Annu. Rev. Psychol. 62, 73–101.

Cohen, J.R., and D’Esposito, M. (2016). The Segregation and Integration of Distinct Brain Networks and Their Relationship to Cognition. J. Neurosci. 36, 12083–12094.

Consortium, T.A.-200 (2012). The ADHD-200 Consortium: A Model to Advance the Translational Potential of Neuroimaging in Clinical Neuroscience. Front. Syst. Neurosci. 6, 62.

Corbetta, M., and Shulman, G.L. (2002). CONTROL OF GOAL-DIRECTED AND STIMULUS-DRIVEN ATTENTION IN THE BRAIN. Nat. Rev. Neurosci. 3, 215–229.

Corbetta, M., Shulman, G.L., Miezin, F.M., and Petersen, S.E. (1995). Superior parietal cortex activation during spatial attention shifts and visual feature conjunction. Science (80-.). 270, 802–805.

Coull, J.T., Vidal, F., Nazarian, B., and Macar, F. (2004). Functional Anatomy of the Attentional Modulation of Time Estimation. Science (80-.). 303, 1506–1508.

Cox, R.W. (1996). AFNI: Software for analysis and visualization of functional magnetic resonance neuroimages. Comput. Biomed. Res. 29, 162–173.

Dan, L., and Yu, J. (1990). Report on Shanghai norms for the Chinese translation of the Wechsler Intelligence Scale for Children - Revised. Psychol. Rep. 67, 531–541.

Deary, I.J., Penke, L., and Johnson, W. (2010). The neuroscience of human intelligence differences. Nat. Rev. Neurosci. 11, 201–211.

DuPaul, G.J., Power, T.J., Anastopoulos, A.D., and Reid, R. (1998). ADHD Rating Scale—IV: Checklists, norms, and clinical interpretation. - PsycNET (Guilford Press, New York).

Engle, R.W. (2002). Working Memory Capacity as Executive Attention. Curr. Dir. Psychol. Sci. 11, 19–23.

Esterman, M., Noonan, S.K., Rosenberg, M., and Degutis, J. (2013). In the zone or zoning out? Tracking behavioral and neural fluctuations during sustained attention. Cereb. Cortex 23, 2712– 2723.

Fan, J., McCandliss, B.D., Fossella, J., Flombaum, J.I., and Posner, M.I. (2005). The activation of attentional networks. Neuroimage 26, 471–479.

Finn, E.S., and Bandettini, P.A. (2020). Movie-watching outperforms rest for functional connectivity-based prediction of behavior. BioRxiv 2020.08.23.263723.

Finn, E.S., Shen, X., Scheinost, D., Rosenberg, M.D., Huang, J., Chun, M.M., Papademetris, X., and Constable, R.T. (2015). Functional connectome fingerprinting: identifying individuals using patterns of brain connectivity. Nat. Neurosci. 18, 1664–1671.

Gao, J.H., Parsons, L.M., Bower, J.M., Xiong, J., Li, J., and Fox, P.T. (1996). Cerebellum implicated in sensory acquisition and discrimination rather than motor control. Science (80-.). 272, 545–547.

Gottwald, B., Mihajlovic, Z., Wilde, B., and Mehdorn, H.M. (2003). Does the cerebellum contribute to specific aspects of attention? Neuropsychologia 41, 1452–1460.

Gratton, C., Kraus, B.T., Greene, D.J., Gordon, E.M., Laumann, T.O., Nelson, S.M., Dosenbach, N.U.F., and Petersen, S.E. (2020). Defining Individual-Specific Functional Neuroanatomy for Precision Psychiatry. Biol. Psychiatry 88, 28–39.

Greene, A.S., Gao, S., Scheinost, D., and Constable, R.T. (2018). Task-induced brain state manipulation improves prediction of individual traits. Nat. Commun. 9, 2807.

Heinrichs, R.W., and Zakzanis, K.K. (1998). Neurocognitive deficit in schizophrenia: A quantitative review of the evidence. Neuropsychology 12, 426–445.

Heinze, H.J., Mangun, G.R., Burchert, W., Hinrichs, H., Scholz, M., Münte, T.F., Gös, A., Scherg, M., Johannes, S., Hundeshagen, H., et al. (1994). Combined spatial and temporal imaging of brain activity during visual selective attention in humans. Nature 372, 543–546.

Hopfinger, J.B., Buonocore, M.H., and Mangun, G.R. (2000). The neural mechanisms of top-down attentional control. Nat. Neurosci. 3, 284–291.

Hsu, W.-T., Rosenberg, M.D., Scheinost, D., Constable, R.T., and Chun, M.M. (2018). Resting-state functional connectivity predicts neuroticism and extraversion in novel individuals. Soc. Cogn. Affect. Neurosci. 13, 224–232.

Huang, L., Mo, L., and Li, Y. (2012). Measuring the interrelations among multiple paradigms of visual attention: An individual differences approach. J. Exp. Psychol. Hum. Percept. Perform. 38, 414–428.

Jiang, R., Calhoun, V.D., Zuo, N., Lin, D., Li, J., Fan, L., Qi, S., Sun, H., Fu, Z., Song, M., et al. (2018). Connectome-based individualized prediction of temperament trait scores. Neuroimage 183, 366–374.

Jiang, R., Zuo, N., Ford, J.M., Qi, S., Zhi, D., Zhuo, C., Xu, Y., Fu, Z., Bustillo, J., Turner, J.A., et al. (2020). Task-induced brain connectivity promotes the detection of individual differences in brain-behavior relationships. Neuroimage 207, 116370.

Kanwisher, N., and Wojciulik, E. (2000). Visual attention: Insights from brain imaging. Nat. Rev. Neurosci. 1, 91–100.

Kucyi, A., Hove, M.J., Esterman, M., Hutchison, R.M., and Valera, E.M. (2017). Dynamic Brain Network Correlates of Spontaneous Fluctuations in Attention. Cereb. Cortex 27, 1831–1840.

Leiner, H.C., Leiner, A.L., and Dow, R.S. (1986). Does the Cerebellum Contribute to Mental Skills? Behav. Neurosci. 100, 443–454.

Levin, H.S., Mattis, S., Ruff, R.M., Eisenberg, H.M., Marshall, L.F., Tabaddor, K., High, W.M., and Frankowski, R.F. (1987). Neurobehavioral outcome following minor head injury: A three-center study. J. Neurosurg. 66, 234–243.

Lin, Q., Rosenberg, M.D., Yoo, K., Hsu, T.W., O’Connell, T.P., and Chun, M.M. (2018). Resting-State Functional Connectivity Predicts Cognitive Impairment Related to Alzheimer’s Disease. Front. Aging Neurosci. 10, 94.

Luck, S.J., and Vogel, E.K. (1997). The capacity of visual working memory for features and conjunctions. Nature 390, 279–284.

Meer, J.N. van der Breakspear, M., Chang, L.J., Sonkusare, S., and Cocchi, L. (2020). Movie viewing elicits rich and reliable brain state dynamics. Nat. Commun. 11, 5004.

Miyake, A., Friedman, N.P., Emerson, M.J., Witzki, A.H., Howerter, A., and Wager, T.D. (2000). The Unity and Diversity of Executive Functions and Their Contributions to Complex “Frontal Lobe” Tasks: A Latent Variable Analysis. Cogn. Psychol. 41, 49–100.

Noble, S., Spann, M.N., Tokoglu, F., Shen, X., Constable, R.T., and Scheinost, D. (2017). Influences on the Test-Retest Reliability of Functional Connectivity MRI and its Relationship with Behavioral Utility. Cereb. Cortex 27, 5415–5429.

Pardo, J. V., Fox, P.T., and Raichle, M.E. (1991). Localization of a human system for sustained attention by positron emission tomography. Nature 349, 61–64.

Pashler, H. (1988). Familiarity and visual change detection. Percept. Psychophys. 44, 369–378.

Petersen, S.E., Fox, P.T., Posner, M.I., Mintun, M., and Raichle, M.E. (1989). Positron emission tomographic studies of the processing of single words. J. Cogn. Neurosci. 1, 153–170.

Pujol, J., Conesa, G., Deus, J., López-Obarrio, L., Isamat, F., and Capdevila, A. (1998). Clinical application of functional magnetic resonance imaging in presurgical identification of the central sulcus. J. Neurosurg. 88, 863–869.

Pylyshyn, Z.W., and Storm, R.W. (1988). Tracking multiple independent targets: evidence for a parallel tracking mechanism. Spat. Vis. 3, 179–197.

Rees, G., Frackowiak, R., and Frith, C. (1997). Two modulatory effects of attention that mediate object categorization in human cortex. Science (80-.). 275, 835–838.

Rosenberg, M., Noonan, S., DeGutis, J., and Esterman, M. (2013). Sustaining visual attention in the face of distraction: a novel gradual-onset continuous performance task. Atten. Percept. Psychophys. 75, 426–439.

Rosenberg, M.D., Finn, E.S., Scheinost, D., Papademetris, X., Shen, X., Constable, R.T., and Chun, M.M. (2016a). A neuromarker of sustained attention from whole-brain functional connectivity. Nat. Neurosci. 19, 165–171.

Rosenberg, M.D., Zhang, S., Hsu, W.-T., Scheinost, D., Finn, E.S., Shen, X., Constable, R.T., Li, C.-S.R., and Chun, M.M. (2016b). Methylphenidate Modulates Functional Network Connectivity to Enhance Attention. J. Neurosci. 36.

Rosenberg, M.D., Finn, E.S., Scheinost, D., Constable, R.T., and Chun, M.M. (2017). Characterizing Attention with Predictive Network Models. Trends Cogn. Sci. 21, 290–302.

Rosenberg, M.D., Hsu, W.-T., Scheinost, D., Todd Constable, R., and Chun, M.M. (2018). Connectome-based Models Predict Separable Components of Attention in Novel Individuals. J. Cogn. Neurosci. 30, 160–173.

Rosenberg, M.D., Scheinost, D., Greene, A.S., Avery, E.W., Kwon, Y.H., Finn, E.S., Ramani, R., Qiu, M., Todd Constable, R., and Chun, M.M. (2020). Functional connectivity predicts changes in attention observed across minutes, days, and months. Proc. Natl. Acad. Sci. U. S. A. 117, 3797–3807.

Satterthwaite, T.D., Elliott, M.A., Ruparel, K., Loughead, J., Prabhakaran, K., Calkins, M.E., Hopson, R., Jackson, C., Keefe, J., Riley, M., et al. (2014). Neuroimaging of the Philadelphia Neurodevelopmental Cohort. Neuroimage 86, 544–553.

Scheinost, D., Noble, S., Horien, C., Greene, A.S., Lake, E.M., Salehi, M., Gao, S., Shen, X., O’Connor, D., Barron, D.S., et al. (2019). Ten simple rules for predictive modeling of individual differences in neuroimaging. Neuroimage 193, 35–45.

Shen, X., Tokoglu, F., Papademetris, X., and Constable, R.T. (2013). Groupwise whole-brain parcellation from resting-state fMRI data for network node identification. Neuroimage 82, 403– 415.

Sprague, T.C., and Serences, J.T. (2013). Attention modulates spatial priority maps in the human occipital, parietal and frontal cortices. Nat. Neurosci. 16, 1879–1887.

Stoodley, C.J. (2012). The cerebellum and cognition: Evidence from functional imaging studies. In Cerebellum, (Springer), pp. 352–365.

Strick, P.L., Dum, R.P., and Fiez, J.A. (2009). Cerebellum and Nonmotor Function. Annu. Rev. Neurosci. 32, 413–434.

Tomasi, D., and Volkow, N.D. (2020). Network connectivity predicts language processing in healthy adults. Hum. Brain Mapp. 41, 3696–3708.

Vanderwal, T., Kelly, C., Eilbott, J., Mayes, L.C., and Castellanos, F.X. (2015). Inscapes: A movie paradigm to improve compliance in functional magnetic resonance imaging. Neuroimage 122, 222–232.

Vanderwal, T., Eilbott, J., Finn, E.S., Craddock, R.C., Turnbull, A., and Castellanos, F.X. (2017). Individual differences in functional connectivity during naturalistic viewing conditions. Neuroimage 157, 521–530.

Wang, J., Ren, Y., Hu, X., Nguyen, V.T., Guo, L., Han, J., and Guo, C.C. (2017). Test-retest reliability of functional connectivity networks during naturalistic fMRI paradigms. Hum. Brain Mapp. 38, 2226–2241.

Weissman, D.H., Roberts, K.C., Visscher, K.M., and Woldorff, M.G. (2006). The neural bases of momentary lapses in attention. Nat. Neurosci. 9, 971–978.

Wimmer, R.D., Schmitt, L.I., Davidson, T.J., Nakajima, M., Deisseroth, K., and Halassa, M.M. (2015). Thalamic control of sensory selection in divided attention. Nature 526, 705–709.

Wojciulik, E., and Kanwisher, N. (1999). The generality of parietal involvement in visual attention. Neuron 23, 747–764.

Woo, C., Chang, L.J., Lindquist, M.A., and Wager, T.D. (2017). Building better biomarkers&: brain models in translational neuroimaging. 20, 365–377.

Wu, E.X.W., Liaw, G.J., Goh, R.Z., Chia, T.T.Y., Chee, A.M.J., Obana, T., Rosenberg, M.D., Yeo, B.T.T., and Asplund, C.L. (2020). Overlapping attentional networks yield divergent behavioral predictions across tasks: Neuromarkers for diffuse and focused attention? Neuroimage 209, 116535.

Yoo, K., Rosenberg, M.D., Hsu, W.T., Zhang, S., Li, C.S.R., Scheinost, D., Constable, R.T., and Chun, M.M. (2018). Connectome-based predictive modeling of attention: Comparing different functional connectivity features and prediction methods across datasets. Neuroimage 167, 11– 22.

Yoo, K., Rosenberg, M.D., Noble, S., Scheinost, D., Constable, R.T., and Chun, M.M. (2019). Multivariate approaches improve the reliability and validity of functional connectivity and prediction of individual behaviors. Neuroimage 197, 212–223.

Yoo, K., Rosenberg, M.D., Kwon, Y.H., Scheinost, D., Todd Constable, R., and Chun, M.M. (2020). A cognitive state transformation model for task-general and task-specific subsystems of the brain connectome. BioRxiv 2020.12.23.424176.

Zhang, H., Hao, S., Lee, A., Eickhoff, S.B., Pecheva, D., Cai, S., Meaney, M., Chong, Y., Broekman, B.F.P., Fortier, M. V., et al. (2020). Do intrinsic brain functional networks predict working memory from childhood to adulthood? Hum. Brain Mapp. hbm.25143.

