## Supplemental Figures and Tables for "A brain-based universal measure of attention: predicting task-general and task-specific attention performance and their underlying neural mechanisms from task and resting state fMRI"

### Supplementary Information

#### Supplementary Tables

| | | Head motion during fMRI scanning | | | | |
|---|---|---|---|---|---|---|
| | | gradCPT | MOT | VSTM | Rest | Movie |
| Performance in tasks | gradCPT | 0.805 | 0.711 | 0.636 | 0.360 | 0.822 |
| | MOT | **$2.31 \times 10^{-4}$** | **$4.29 \times 10^{-4}$** | **$3.23 \times 10^{-3}$** | **$8.43 \times 10^{-4}$** | **$1.37 \times 10^{-3}$** |
| | VSTM | 0.211 | 0.091 | 0.306 | 0.075 | 0.227 |

**Supplementary Table S1**. Correlation between head motion (mean framewise displacement) and behavioral performance in tasks. Parametric p-values of Pearson's correlation are reported. Significant correlations under uncorrected $p<0.05$ are shown in **bold**. GradCPT: gradual-onset continuous performance task, MOT: multiple object tracking, and VSTM: visual short-term memory.

| fMRI data | | gradCPT | MOT | VSTM | Rest | Movie |
|---|---|---|---|---|---|---|
| Predictive connectivity # | Positive network | 82 | 110 | 63 | 15 | 32 |
| | Negative network | 97 | 113 | 70 | 99 | 118 |

**Supplementary Table S2**. The numbers of features (i.e., predictive functional connections) in CPMs of the general attention component with three tasks (gradCPT [gradual-onset continuous performance task], MOT [multiple object tracking], and VSTM [visual short-term memory]), rest, and movie fMRI data.

| Network | MF | FPN | DMN | Mot | VI | VII | VAs | SN | Subc | Cbl |
|---|---|---|---|---|---|---|---|---|---|---|
| Nodes # | 27 | 28 | 18 | 44 | 18 | 9 | 17 | 29 | 29 | 27 |

**Supplementary Table S3**. The number of nodes in ten networks. MF: medial-frontal network, FPN: frontoparietal network, DMN: default mode network, Mot: motor, VI: visual I, VII: visual II, VAs: visual associations, SN: salience network, Subc: subcortex, Cbl: cerebellum.

| | | PC 1 | PC 2 | PC 3 |
|---|---|---|---|---|
| Variance explained (%) | | 62.4 | 21.2 | 16.4 |
| Correlation | gradCPT | 0.776 | -0.530 | -0.341 |
| | MOT | 0.763 | 0.593 | -0.259 |
| | VSTM | 0.829 | -0.049 | 0.558 |

**Supplementary Table S4**. Principal components of three attentional behaviors. GradCPT: gradual-onset continuous performance task, MOT: multiple object tracking, and VSTM: visual short-term memory.

### Supplementary Figures

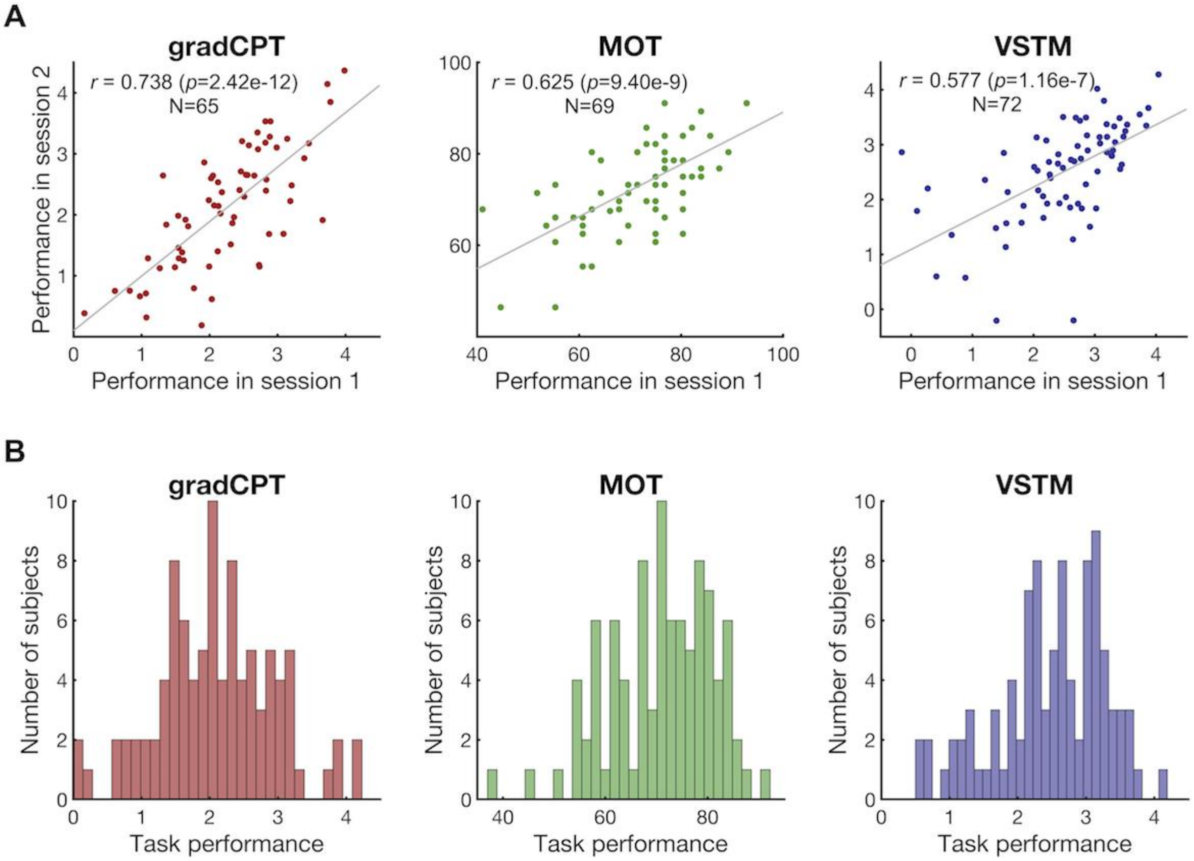


**Supplementary Figure S1**. **A**. A reliability of task performance measures for each task. The significant correlation indicates that individual behaviors were reliably measured. **B**. Distribution of individual performances averaged between two sessions for three attention tasks. GradCPT: gradual-onset continuous performance task, MOT: multiple object tracking, and VSTM: visual short-term memory.

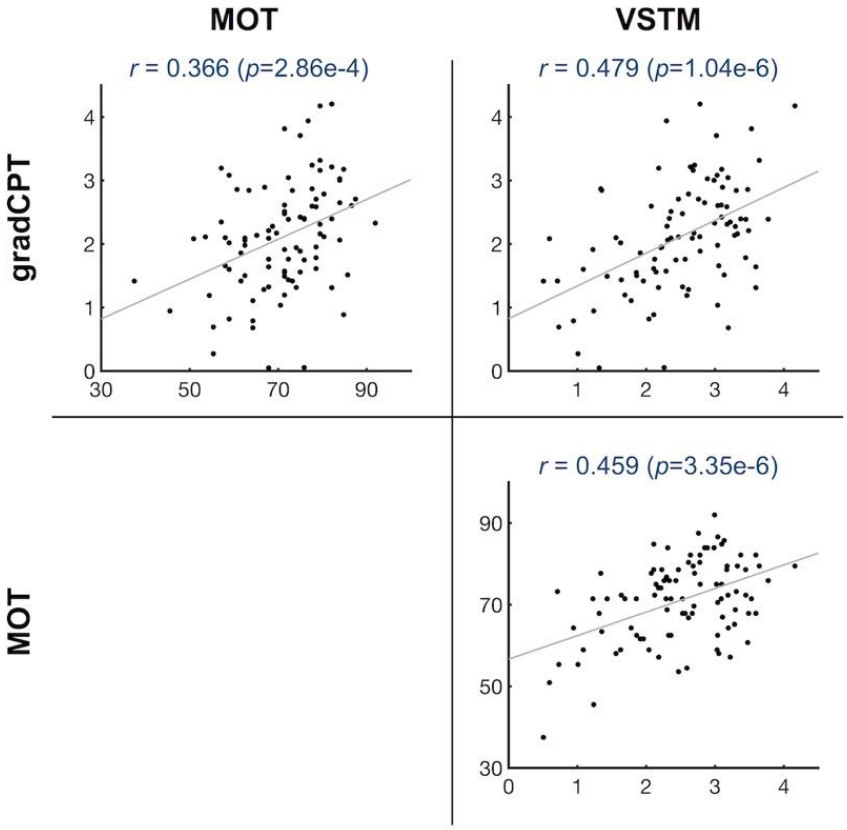


**Supplementary Figure S2.** A similarity of individual behaviors between different tasks. To assess the similarity, we estimated correlation of individual performances between attention tasks. Individual behaviors were significantly correlated between every pair of tasks. GradCPT: gradual-onset continuous performance task, MOT: multiple object tracking, and VSTM: visual short-term memory.

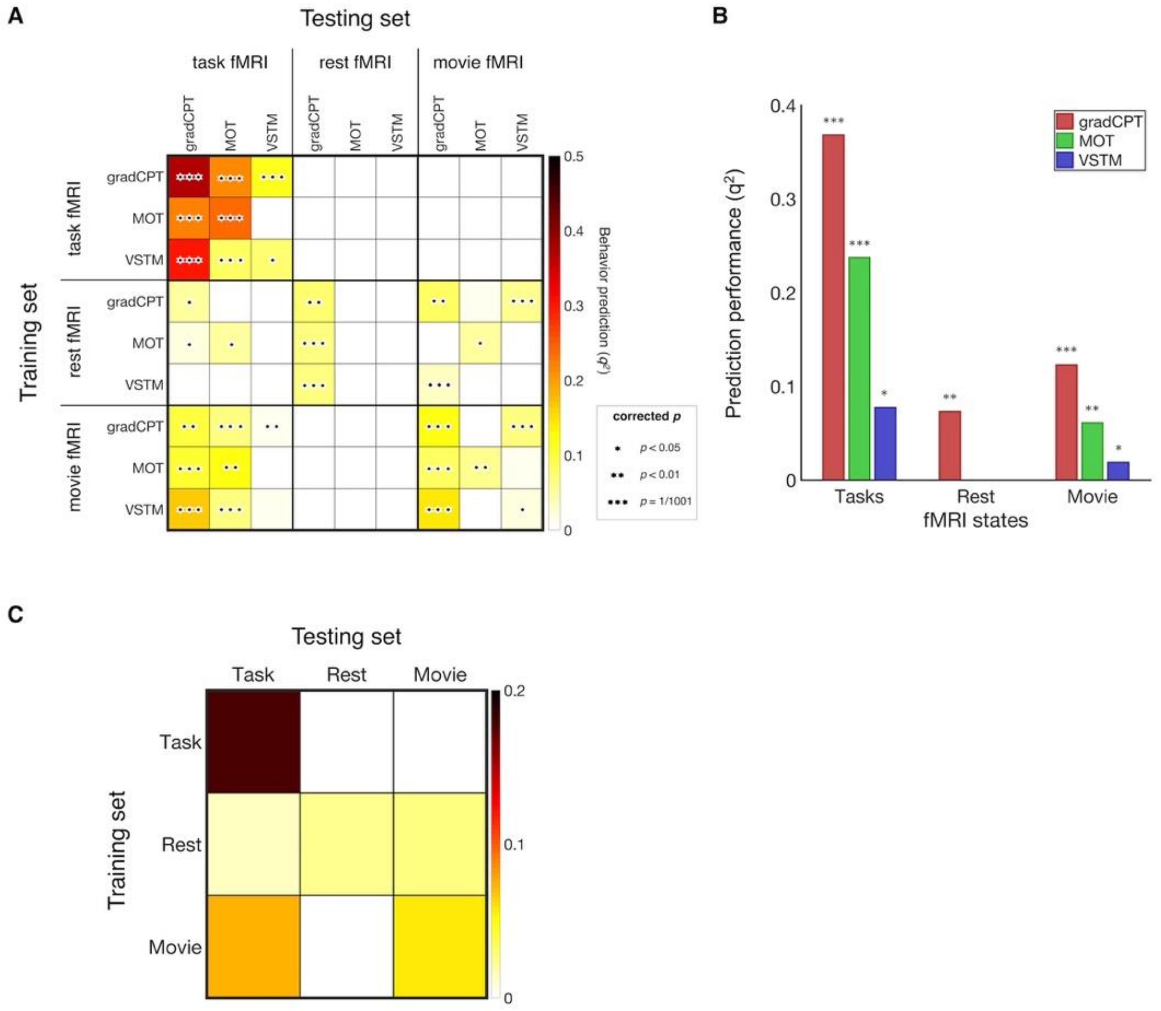


**Supplementary Figure S3**. Prediction accuracy of nine original CPMs measured using prediction $q^2$ that is based on normalized mean square error. **A**. Cross-prediction results of the original models (Figure 2) across all cognitive states and attention tasks. P-values corrected for multiple tests (three correlations in task fMRI, three in rest fMRI, and three in movie fMRI) were obtained using 1,000 permutation ($***$: $p$=1/1001; $**$: $p$<0.01; $*$: $p$<0.05). GradCPT: gradual-onset continuous performance task, MOT: multiple object tracking, and VSTM: visual short-term memory. **B**. Within-task prediction accuracy of nine CPM models (Figure 1 and 2). This result corresponds to the nine diagonal elements in A. **C**. This panel represents the averaged performance within each fMRI type (performing an attention-task, resting state, or movie-watching) in A. The CPMs trained on task fMRI made the most accurate predictions, but did not generalize to different types of fMRI. In contrast, the CPMs trained on rest fMRI made the least accurate predictions, but generalized to all other types of fMRI, movie and task. The CPMs trained on movie fMRI generalized only to task fMRI.

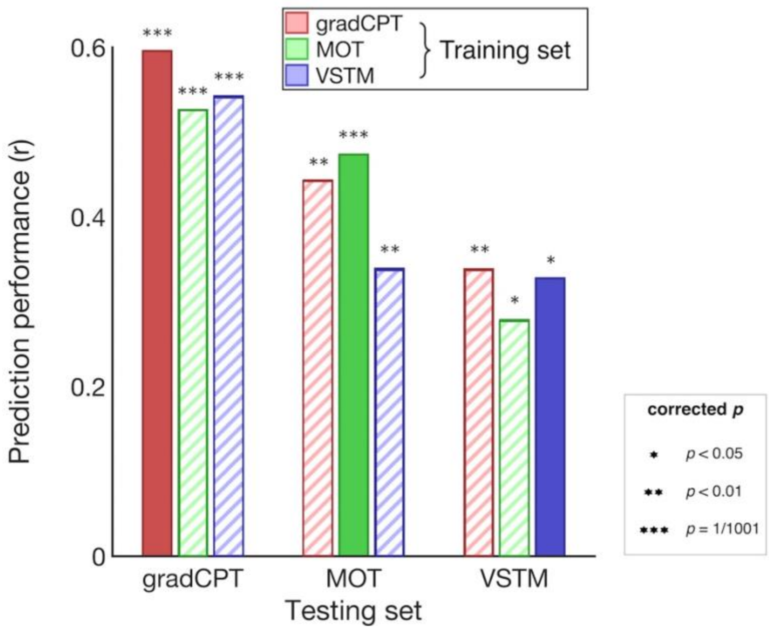


**Supplementary Figure S4.** Task-to-task connectome-based prediction results. Dark filled color represents within-task prediction, and light color with stripes represents cross-task prediction. P-values corrected for multiple tests (three within-task correlation and six cross-task correlation) were obtained using 1,000 permutation (∗∗∗: $p$=1/1001; ∗∗: $p$<0.01; ∗: $p$<0.05). GradCPT: gradual-onset continuous performance task, MOT: multiple object tracking, and VSTM: visual short-term memory.

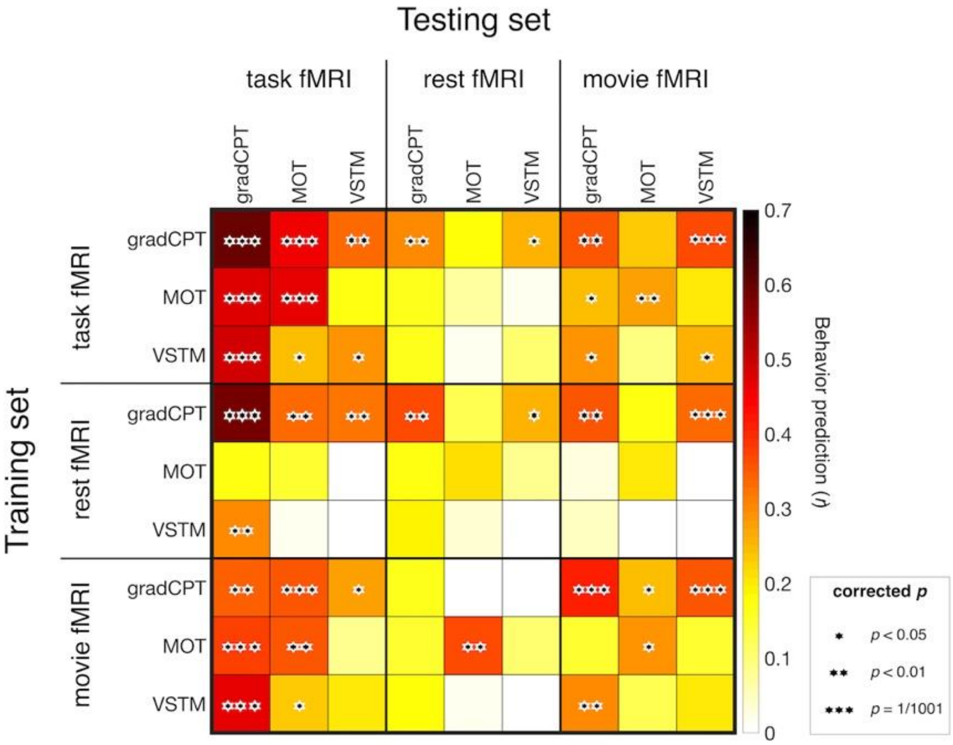


**Supplementary Figure S5**. Cross-prediction results with only task-specific connectivity ($*$: $p$<0.05, $**$: $p$<0.01, and $***$: $p$=1/1,001, corrected using permutation). GradCPT: gradual-onset continuous performance task, MOT: multiple object tracking, and VSTM: visual short-term memory.

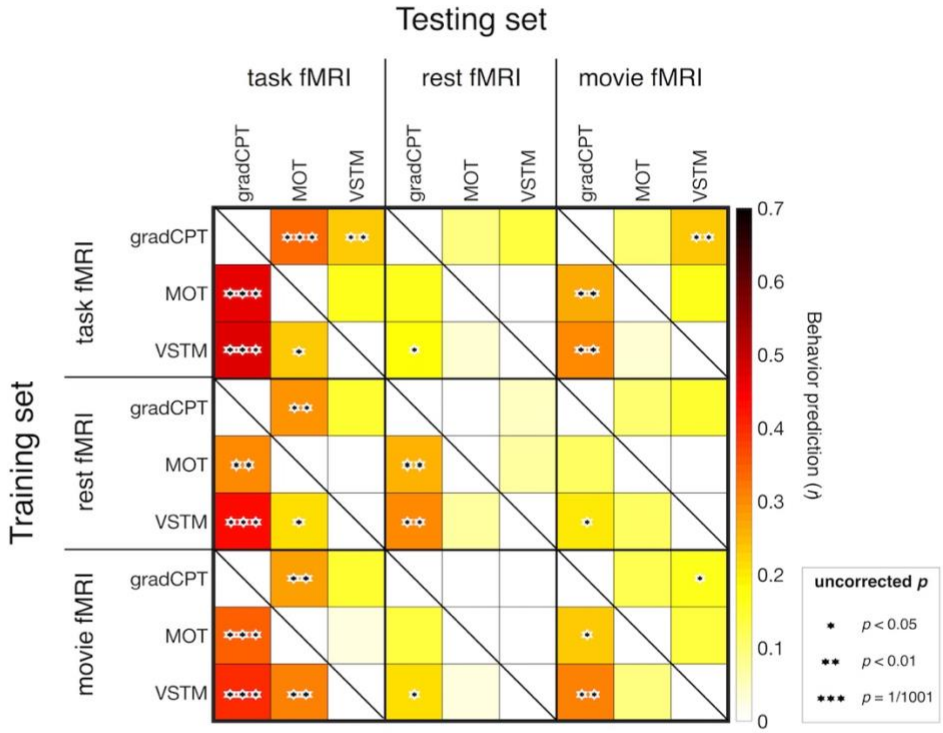


**Supplementary Figure S6.** Cross-prediction results across all cognitive states and attention tasks (∗: $p$<0.05, ∗∗: $p$<0.01, and ∗∗∗: $p$=1/1,001, uncorrected). Performance was assessed by partial correlation to control for correlated behaviors between tasks. GradCPT: gradual-onset continuous performance task, MOT: multiple object tracking, and VSTM: visual short-term memory.

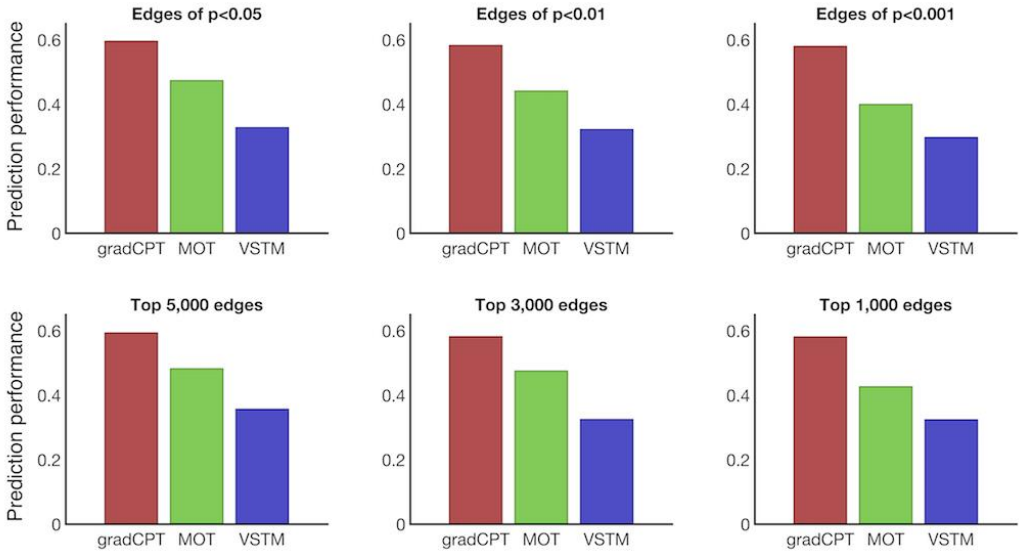


**Supplementary Figure S7**. Controlling for the number of feature edges in CPM. This result confirms that the model prediction is not driven by the number of predictive connectivity features. GradCPT: gradual-onset continuous performance task, MOT: multiple object tracking, and VSTM: visual short-term memory.

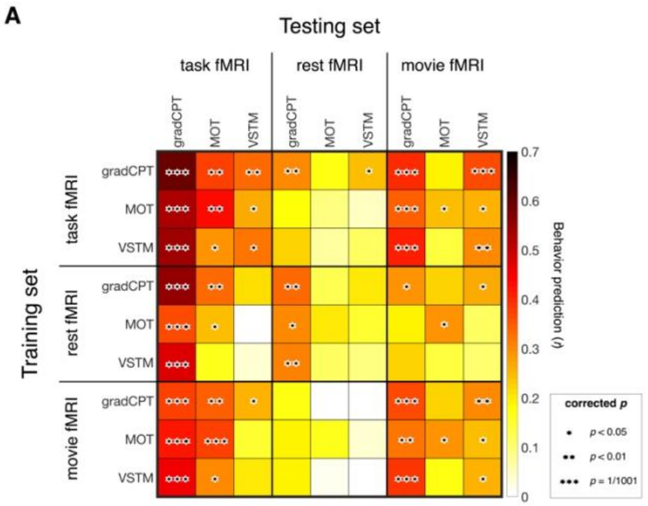


**Supplementary Figure S8**. **A**. Cross-prediction results across all cognitive states and attention tasks with controlling for the head motion measured by mean framewise displacement during fMRI scanning (∗: $p$<0.05, ∗∗: $p$<0.01, and ∗∗∗: $p$=1/1,001, corrected). Performance was assessed by partial correlation to control for head motion. **B**. Decrease in estimates of prediction accuracy after controlling for the head motion compared to the result in **Figure 3**. No significant decrease was observed at $p$<0.05. GradCPT: gradual-onset continuous performance task, MOT: multiple object tracking, and VSTM: visual short-term memory.

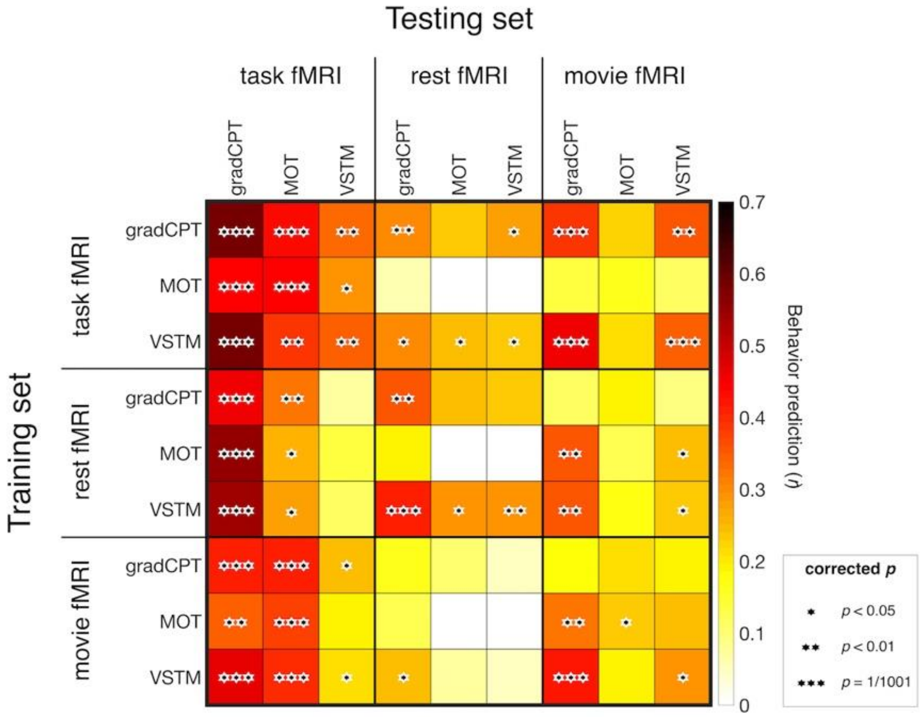


**Supplementary Figure S9**. Cross-prediction results with all task-related edges ($*$: *p*<0.05, $**$: *p*<0.01, and $***$ : *p*=1/1,001, corrected using permutation). The models with task fMRI successfully generalized to different attention tasks (*the top left 3 by 3 subpart*). GradCPT: gradual-onset continuous performance task, MOT: multiple object tracking, and VSTM: visual short-term memory.

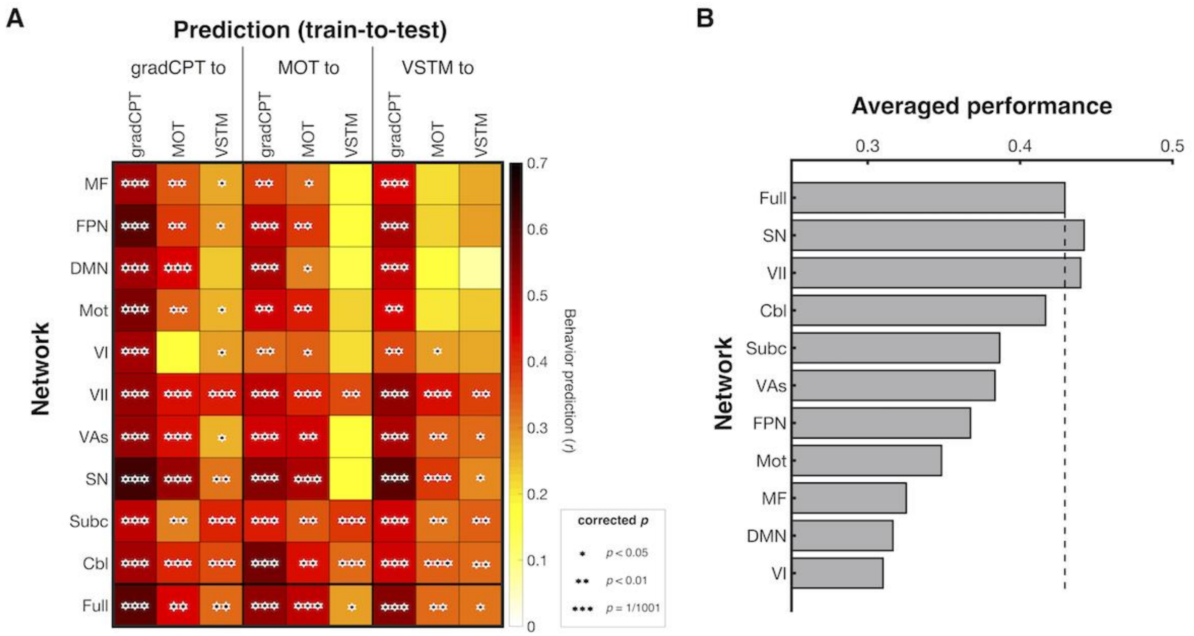


**Supplementary Figure S10.** Cross-prediction using between-network connectivity ($*$: $p$<0.05, $*$: $p$<0.01, and $***$: $p$=1/1,001, corrected using permutation). **A**. Rows represent each canonical network and columns represent combinations of training and testing data. Models' prediction accuracies were assessed by correlating model-predicted and observed behavioral scores. *P* value was obtained using 1,000 permutations and corrected for multiple tests. **B**. The average prediction performance of each network. This was obtained by averaging nine prediction cases in each row in **A**. GradCPT: gradual-onset continuous performance task, MOT: multiple object tracking, and VSTM: visual short-term memory. MF: medial-frontal network, FPN: frontoparietal network, DMN: default mode network, Mot: motor, VI: visual I, VII: visual II, VAs: visual associations, SN: salience network, Subc: subcortex, Cbl: cerebellum.

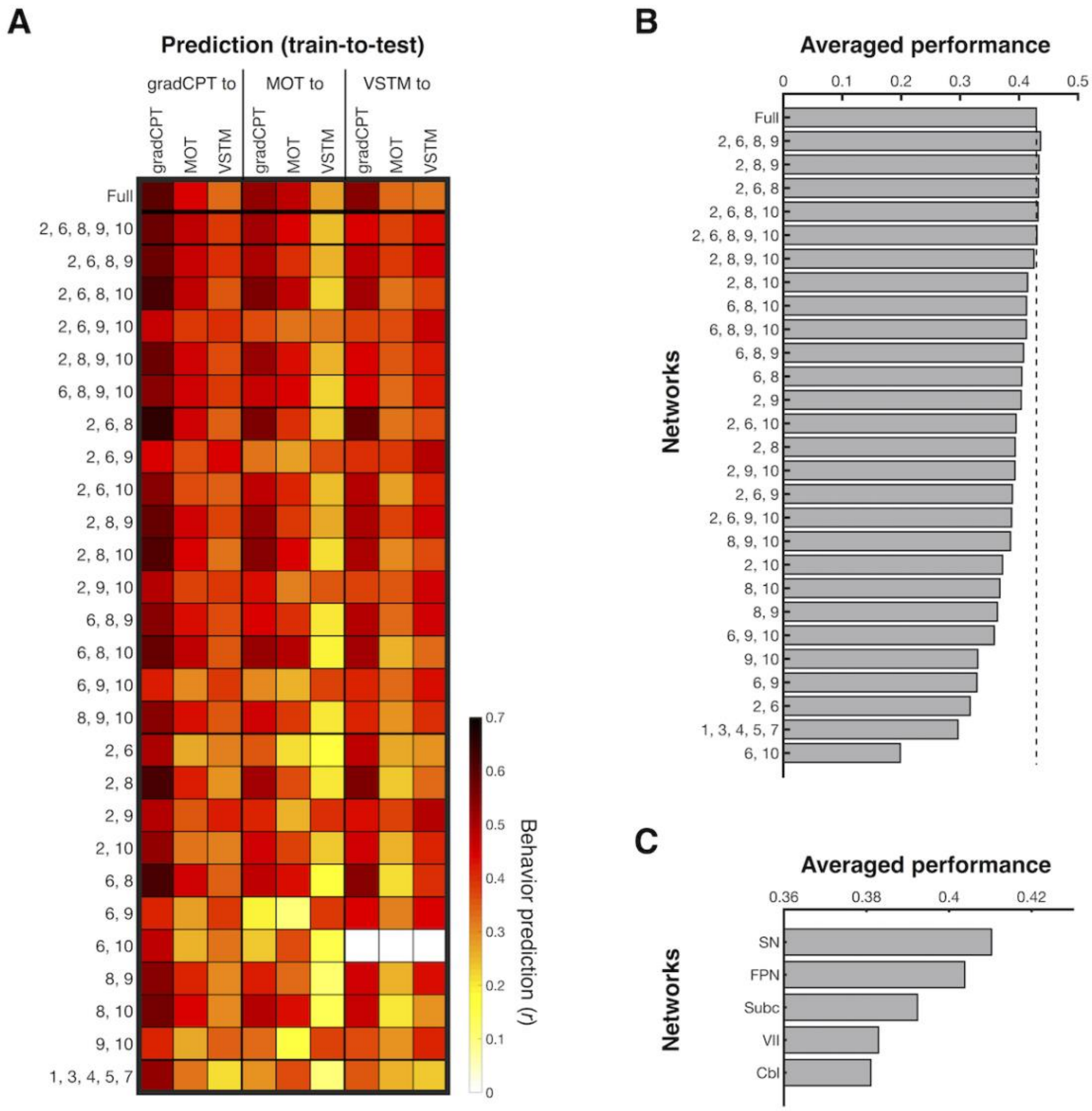


**Supplementary Figure S11**. Cross-prediction using connectivity between the frontoparietal (2), visual II (6), salience (8), subcortical (9), cerebellum (10) networks. Prediction of a model using connectivity between the medial-frontal (1), default mode (3), motor (4), visual I (5), visual association (7) networks was also obtained as a control. **A**. Rows represent networks (indicated by numbers) used in each CPM model, and columns represent combinations of training and testing data. Models' prediction accuracies were assessed by correlating model-predicted and observed behavioral scores. **B**. The average prediction performance of each model. This was obtained by averaging nine prediction cases in each row in **A**. **C**. The average prediction performance of each network. This was obtained for each network by averaging prediction performances of models (tested in **A**) that includes the network. GradCPT: gradual-onset continuous performance task, MOT: multiple object tracking, and VSTM: visual short-term memory.

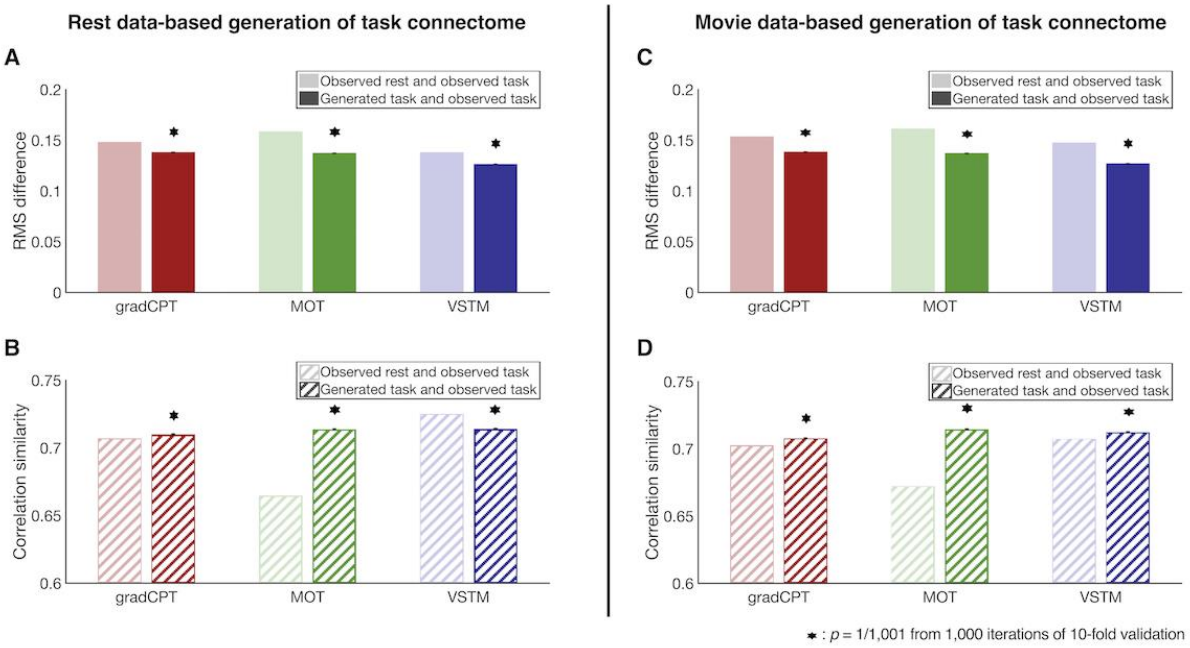


**Supplementary Figure S12**. Similarity between C2C model-generated task connectomes. Error bar represents standard deviation from 1,000 iterations of the mean value across 10-fold cross validation. **A** and **C** represent root mean square (RMS) difference between two connectomes. Hence, the lower difference of the generated connectome indicates that a C2C model accurately generates the target task connectome from the rest connectome. **B** and **D** represent a spatial similarity between two connectomes assessed by correlation. In this case, the higher similarity of the generated connectome indicates that a C2C model accurately generates the target task connectome from the rest connectome. GradCPT: gradual-onset continuous performance task, MOT: multiple object tracking, and VSTM: visual short-term memory.

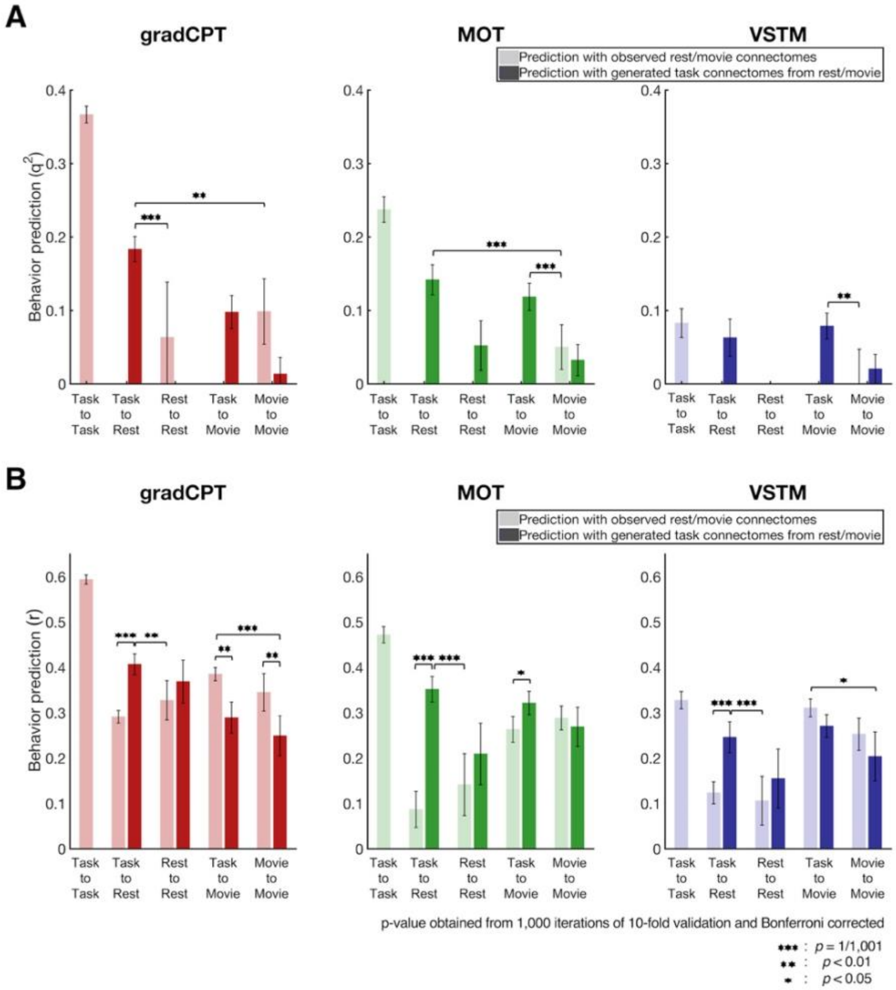


**Supplementary Figure S13**. Prediction of individual behaviors with different approaches. **A**. Prediction performance was assessed by prediction $q^2$ and negative values were set to zero (i.e., task-to-rest prediction with the empirical rest connectome in all three tasks). **B**. Prediction performance was assessed with correlation (*r*). Darker bars represent the behavioral prediction with task connectomes generated by C2C modeling. A darker bar in task-to-rest represents the behavior prediction of a model trained using empirical task connectome and predicted with task connectome generated from empirical rest connectome. A darker bar in rest-to-rest represents the prediction of a model trained using empirical rest connectome and predicted with task connectome generated from rest connectome. A darker bar in task-to-move represents the behavior prediction of a model trained using empirical task connectome and predicted with task connectome generated from empirical movie connectome. A darker bar in movie-to-movie represents the prediction of a model trained using empirical movie connectome and predicted with task connectome generated from movie connectome. The model-generated task connectome from rest data better predicted individual behaviors than empirical rest connectome in all three attention tasks. Prediction performance was assessed by correlating observed and predicted behavioral scores. Error bars represent standard deviation across 1,000 iterations. GradCPT: gradual-onset continuous performance task, MOT: multiple object tracking, and VSTM: visual short-term memory.

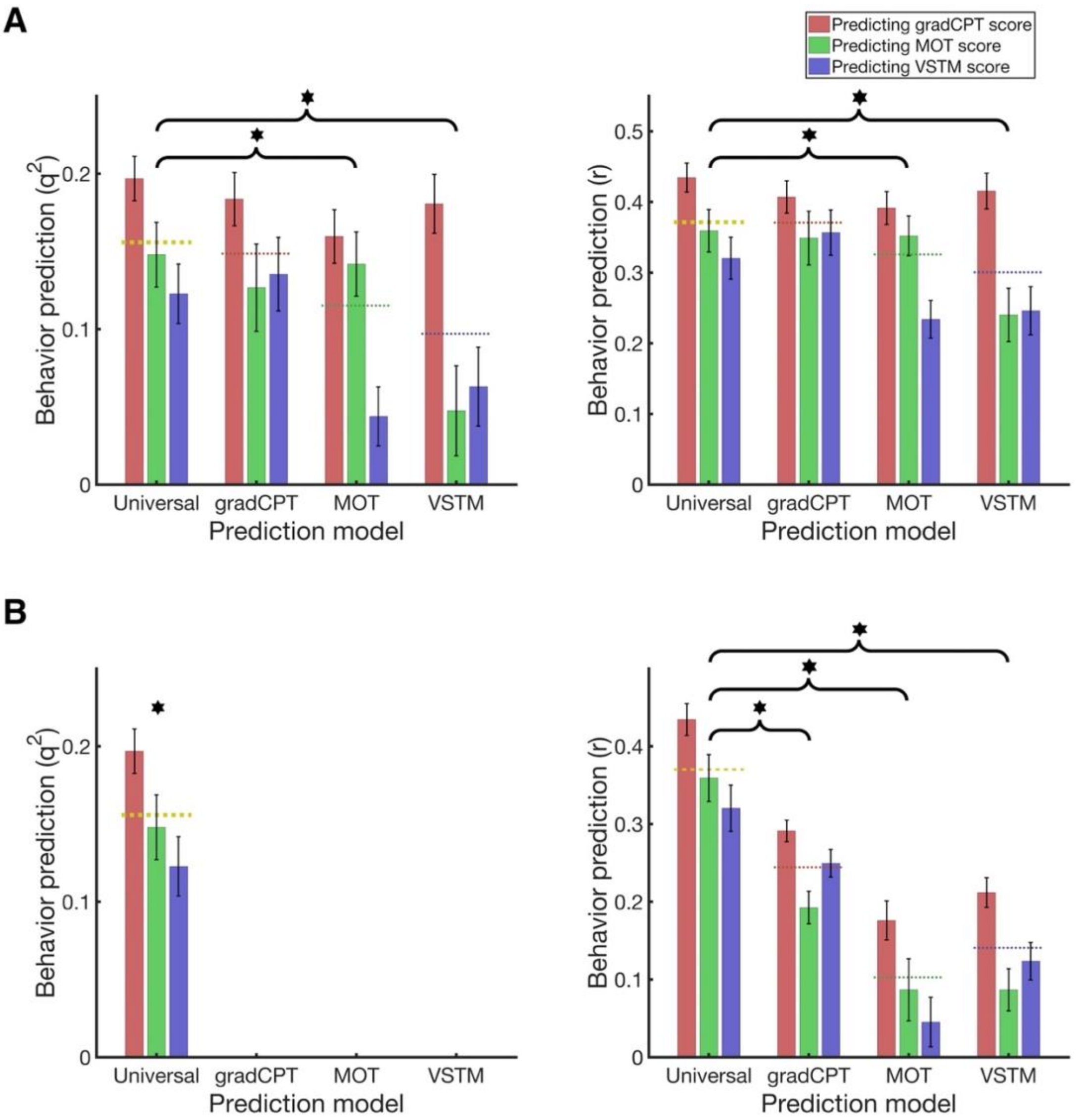


**Supplementary Figure S14**. Behavior prediction by the universal attention model. **A**. Each task name in *x*-axis represents a single task-based model combined with a C2C model implemented. The left panel corresponds to Figure 8. The universal attention model and three CPM models predict individual behaviors from a resting-state connectome. Three colored bars from each model show the model's performance in predicting each of three task behaviors. Dotted lines represent the model performance averaged across three tasks. Error bars represent standard deviation across 1,000 iterations. ∗: The universal model prediction, on average (by dotted lines), significantly better predict behaviors than CPM models at $p<0.05$ from 1,000 iterations. **B**. Each task name in *x*-axis represents a single task-based model without a C2C model implemented. $q^2=0$ in three task-based models (*left*). GradCPT: gradual-onset continuous performance task, MOT: multiple object tracking, and VSTM: visual short-term memory.

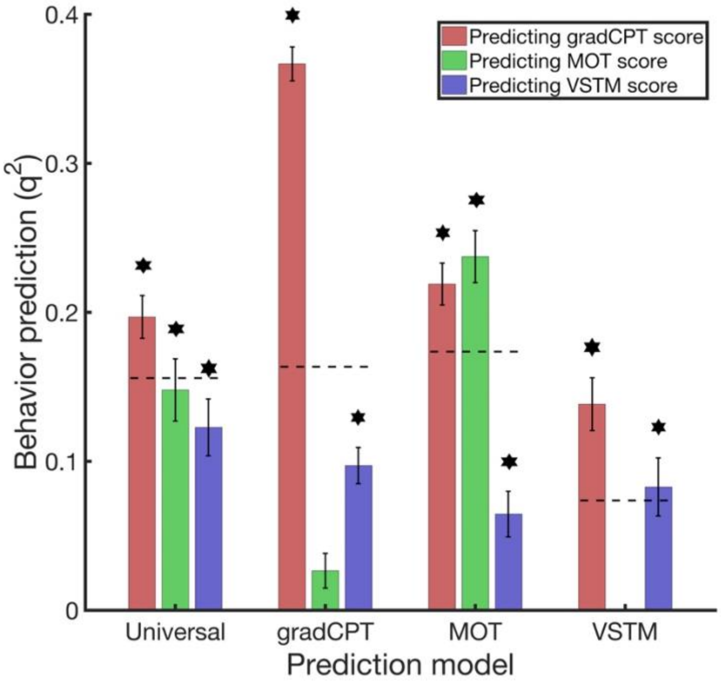


**Supplementary Figure S15**. Behavior prediction by the universal attention model in comparison to the original three task-based CPMs. Each task name in *x*-axis represents a single task-based model. The universal attention model predicts individual behaviors from a resting-state connectome, whereas task-based models predict them from a corresponding task connectome. Three colored bars from each model show the model's performance in predicting each of three task behaviors. Black dotted lines represent the model performance averaged across three tasks. Error bars represent standard deviation across 1,000 iterations. $*$: A model prediction is considered successful if the prediction performance $q^2$ from all 1,000 iterations was above zero. GradCPT: gradual-onset continuous performance task, MOT: multiple object tracking, and VSTM: visual short-term memory.

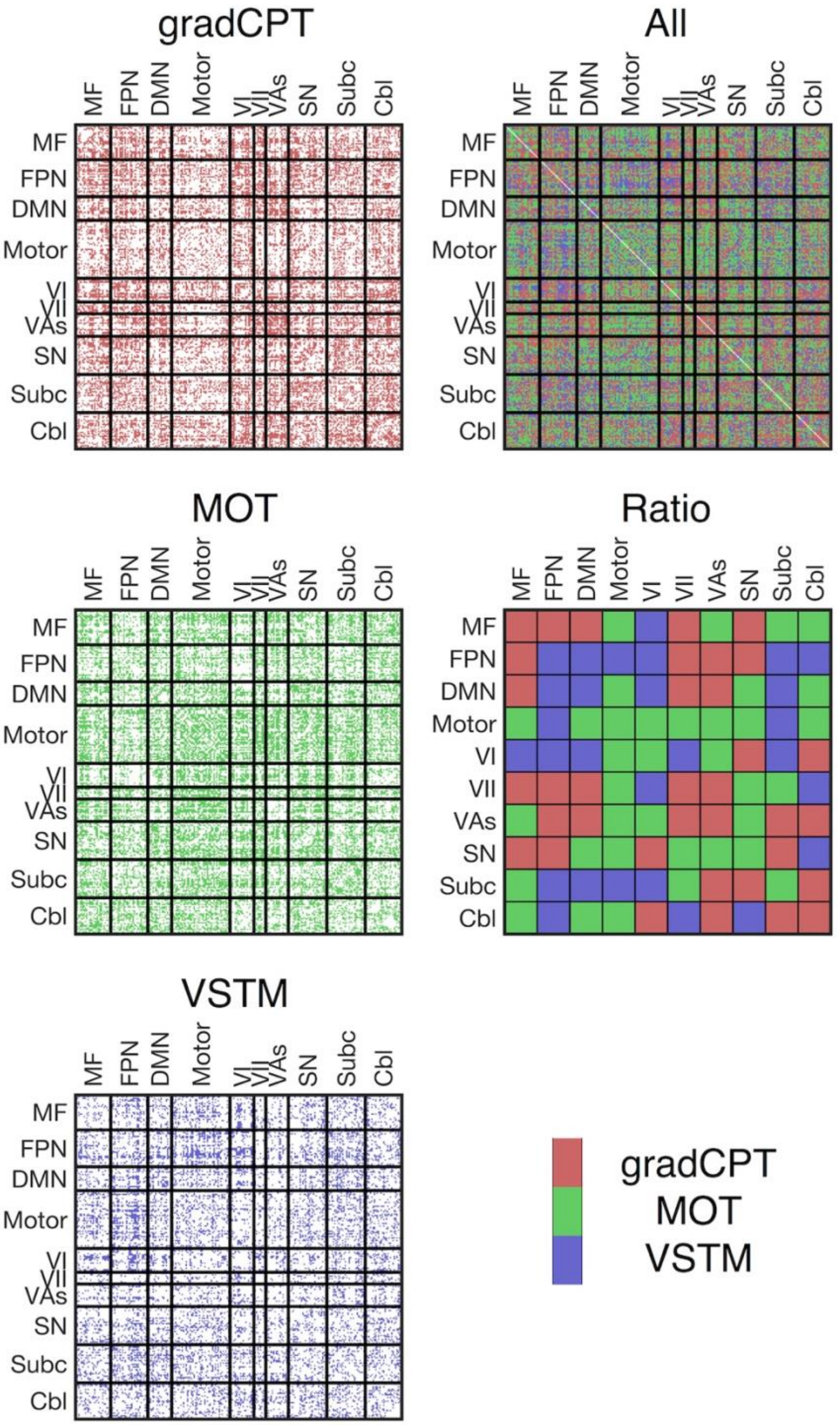


**Supplementary Figure S16**. The universal attention connectome lookup table. Out of a total 30,135 edges, 10,885 (36.1%) edges were pulled from gradCPT, 12,542 (41.6%) edges were from MOT, and 6,708 (22.3%) were from VSTM. The *Ratio* map was obtained based on *All* map. In each within- or between-network element in Ratio, the number of edges in the element for each task was counted and normalized by the total number of edges of each task. A task with the highest normalized value was assigned.

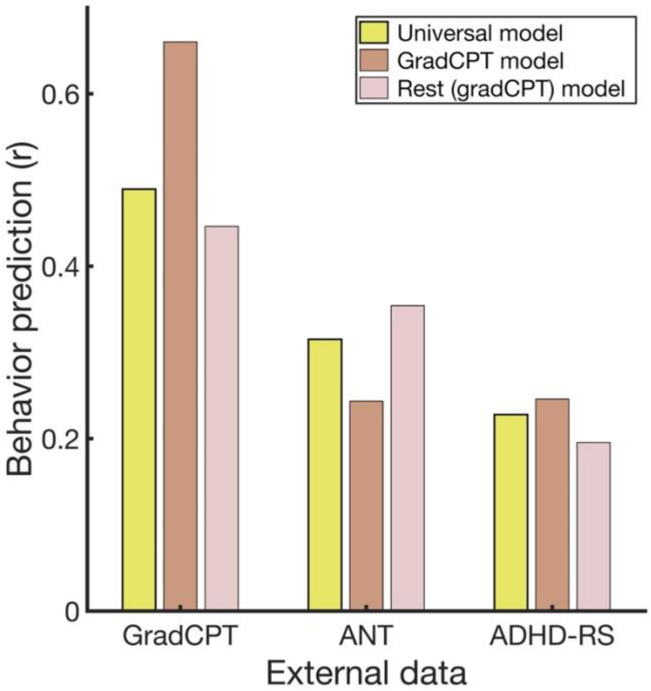


**Supplementary Figure S17**. Generalizability of the universal attention model in three independent datasets. Prediction performance was assessed by correlation *r*. The universal model (yellow) and two CPMs successfully captured individual differences in the external datasets.
